# Nonlinear Dynamics of Left Ventricular Mass Remodeling in Chagas Cardiomyopathy

**DOI:** 10.64898/2026.04.16.719099

**Authors:** Paulo Roberto Benchimol-Barbosa, Ana Carolina L. Benchimol-Barbosa, Claudio Gonçalves Carvalhaes, Bharat K. Kantharia

## Abstract

Left ventricular (LV) mass remodeling in chronic Chagas cardiomyopathy (CCC) is progressive, but whether its dynamics follow nonlinear patterns linked to all-cause mortality remains unexplored. In this SEARCH-Rio substudy, 50 outpatients were followed for 10 years (40 survivors, 10 nonsurvivors). Serial echocardiography provided paired LV mass measurements fitted to the logistic equation x’ = α·x·(1−γ·x), yielding α = 3.952 ± 0.096 and γ = 1.267 ± 0.012, above the complexity threshold (α > 3.57). Lyapunov exponents (LE), computed from consecutive inter patient derivatives, were positive in survivors (+0.349 ± 0.342) and negative in nonsurvivors (−0.459 ± 0.377; p < 0.001), indicating preserved versus lost dynamical complexity, respectively. The fixed-point equilibrium (≈ 179 g) is mathematically unstable for α = 3.952, consistent, but not confirmatory, with sensitive dependence on initial conditions. Complex nonlinear dynamics were independently confirmed at the intra-patient level in a subgroup with extended longitudinal follow-up beyond the initial 10-year period. A Firth penalized regression risk score based on ejection fraction < 51.7% and maximum heart rate < 109 bpm (optimism-corrected AUC = 0.959, bootstrap B = 1,000) showed monotonic mortality gradient and progressively lower LE across strata (Spearman ρ = −0.381, p = 0.003). These findings support nonlinear LV mass remodeling associated with all-cause mortality over the same observation window in CCC; replication in larger cohorts is warranted.

## Introduction

Chronic Chagas cardiomyopathy (CCC), caused by Trypanosoma cruzi infection, remains a leading cause of heart failure and sudden cardiovascular events in Latin America, affecting approximately 6–8 million individuals [1,2]. Left ventricular (LV) remodeling, characterized by progressive changes in the ventricular mass, geometry, and function, is a hallmark of CCC progression [3,4].

The nonlinear dynamical systems theory provides a framework for understanding complex biological processes. Poon and Merrill [5] demonstrated that healthy hearts exhibit deterministic complex nonlinear dynamics in beat-to-beat intervals, with loss of complexity in congestive heart failure (CHF), establishing a paradigm in which the loss of physiological complexity signals disease progression [6]. Wu et al. [7] showed that complex nonlinear heart rate variability (HRV) signatures linked to respiratory sinus arrhythmia distinguish health from CHF. Poon and Barahona [8] demonstrated that the noise limit, which is closely related to the Lyapunov exponent (LE), can be titrated to distinguish deterministic complex dynamics from noise in cardiovascular time series, with implications for mortality prediction in patients with CHF. Arzeno et al. [9] further showed that the loss of nonlinear complexity is the single best predictor of mortality in mild to moderate CHF, outperforming traditional linear HRV indices. The LE quantifies this divergence rate: positive values indicate preserved complexity (small differences amplify over time), whereas negative values indicate convergence (loss of complexity).

Recently, the nonlinear characterization of Chagas disease has been explored using entropy-based HRV analysis. Defeo et al. [10] revealed alterations in heart rate fluctuations during Chagas disease progression. Cornejo et al. [11] and Valdez et al. [12] applied deep learning with permutation entropy to stratify Chagas patients. However, all prior studies have focused on beat-to-beat dynamics. No study has investigated whether structural cardiac remodeling in CCC, measured over years using echocardiography, follows a nonlinear dynamic pattern.

We hypothesized that long-term LV mass remodeling in CCC follows a deterministic nonlinear pattern described by the logistic equation [13,14], and that loss of dynamic complexity is associated with contemporaneous mortality during the period captured by echocardiographic assessments.

## Results

### Baseline Characteristics and Univariate Predictors

Table 1 presents the baseline characteristics of all 50 patients and the univariate comparison between survivors (n=40) and nonsurvivors (n=10) at the 10-year follow-up. Nonsurvivors had shorter follow-up (50.7 ± 28.0 months) than survivors (107.8 ± 36.2 months, Mann-Whitney p<0.001), and were characterized by more advanced disease: higher NYHA class (40 % vs. 0 % class II, Fisher p<0.001), higher Los Andes stage (80 % stage III vs. 10 %, p<0.001), and reduced left ventricular function (median EF 39.1 % vs. 75.3 %, Mann-Whitney p<0.001). Holter monitoring revealed a significantly higher ventricular ectopy burden in nonsurvivors, including PVCs/day (4,929 vs. 56, p=0.002), paired PVCs/day (93 vs. 0, p<0.001), and SVE runs/day (552 vs. 21, p=0.008). NSVT was also significantly more frequent in nonsurvivors (60 % vs. 15 %, p=0.007). Echocardiographic parameters showed larger LV dimensions (LVESD 4.8 vs. 2.9 cm, p<0.001), greater LV mass (210 vs. 141 g, p=0.015), and more frequent mitral regurgitation (90 % vs. 42 %, p=0.011).

**Table 1.**
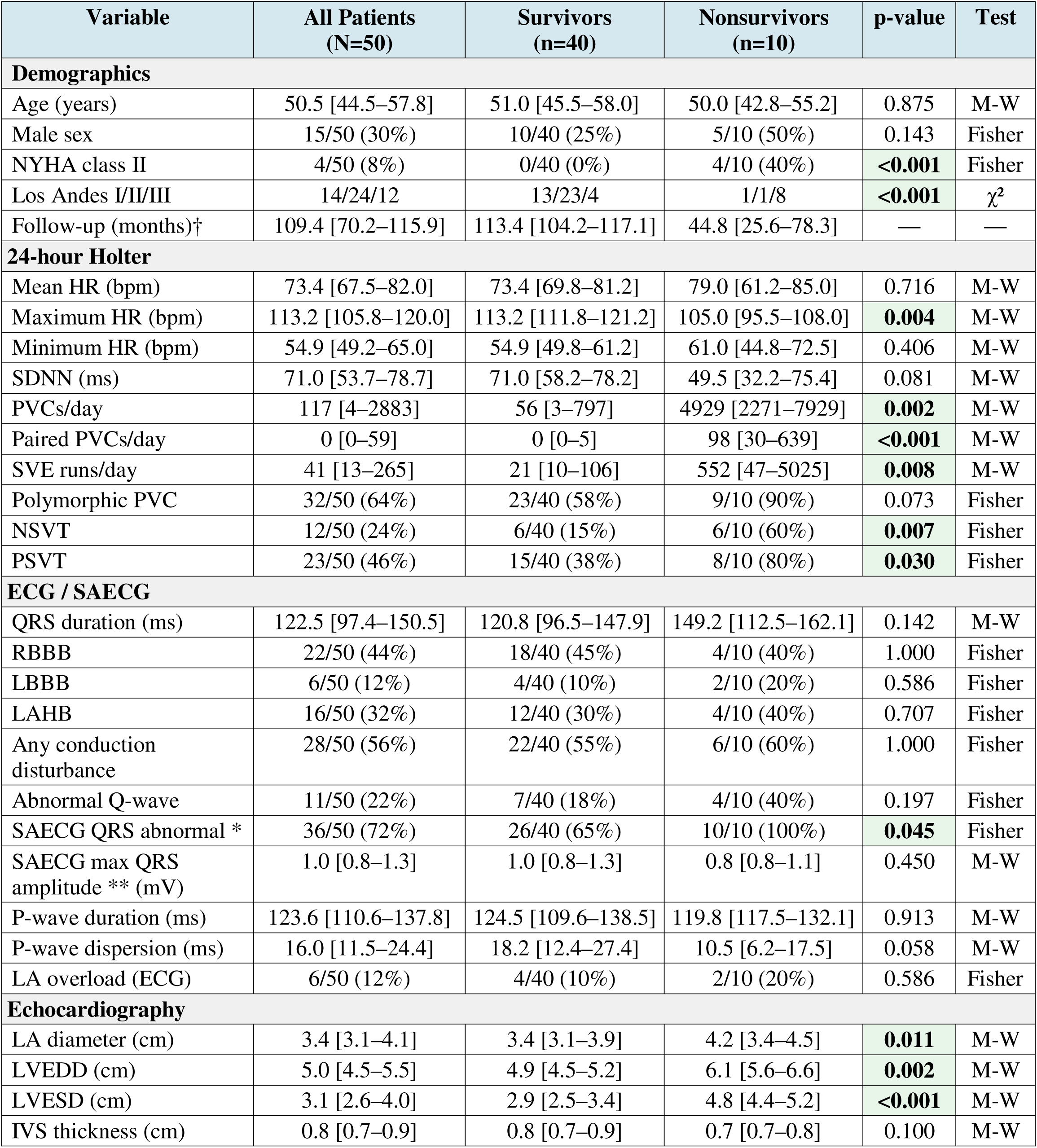
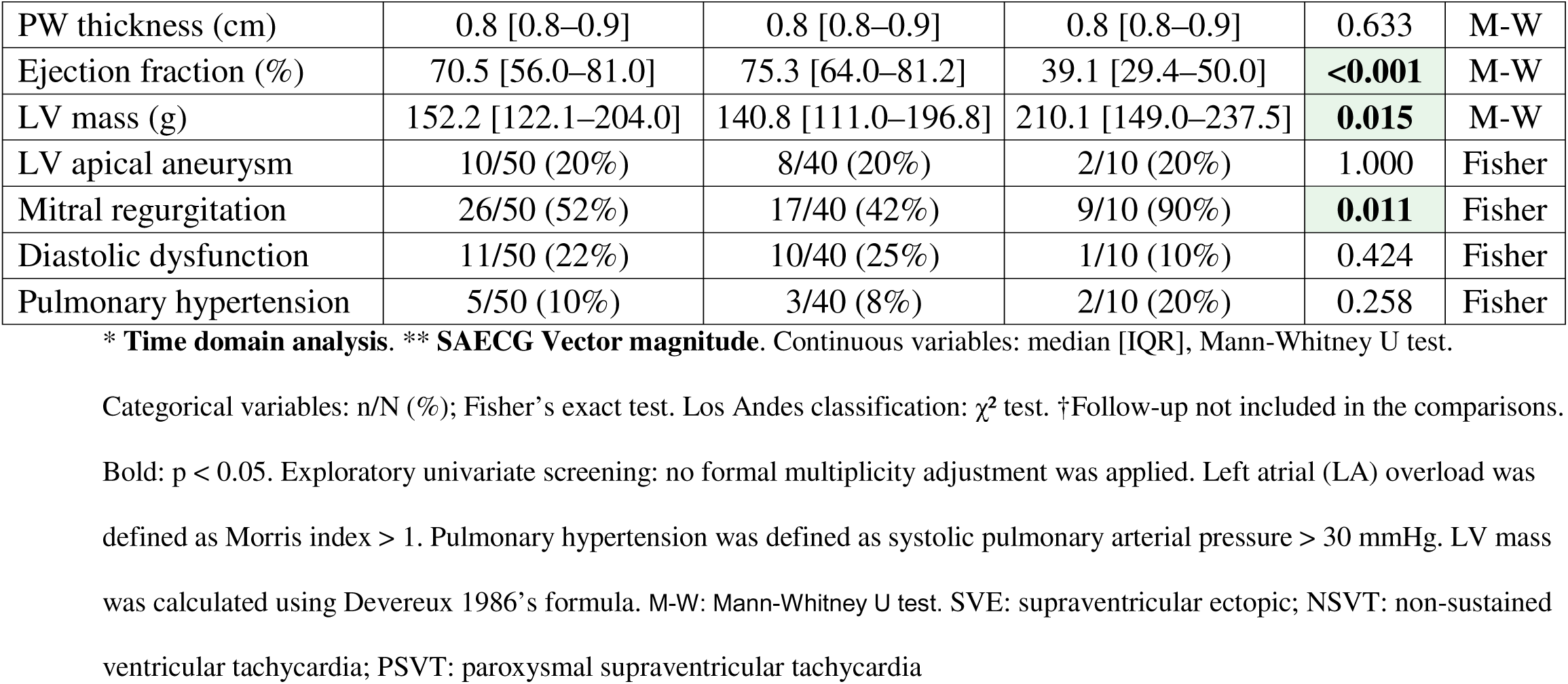
Baseline characteristics and univariate predictors of death within 10 years (10-year follow-up).

#### Logistic Map of LV Mass Remodeling

LV mass remodeling fitted the logistic equation (Figure 1) with α = 3.952 ± 0.096 (CI 3.75 to 4.14), γ = 1.267 ± 0.012 (CI 1.25 to 1.30), and zero convergence failures in 1,000 bootstraps. The fit yielded R² = 98.6 %; given only six grouped points and two parameters (df = 4), this is interpreted as internal goodness-of-fit. The value of α (3.952) places the system above the complexity threshold (α > 3.57), consistent with a complex nonlinear dynamical regime, conditional on the logistic equation being an adequate descriptor.

**Figure 1 legend.**
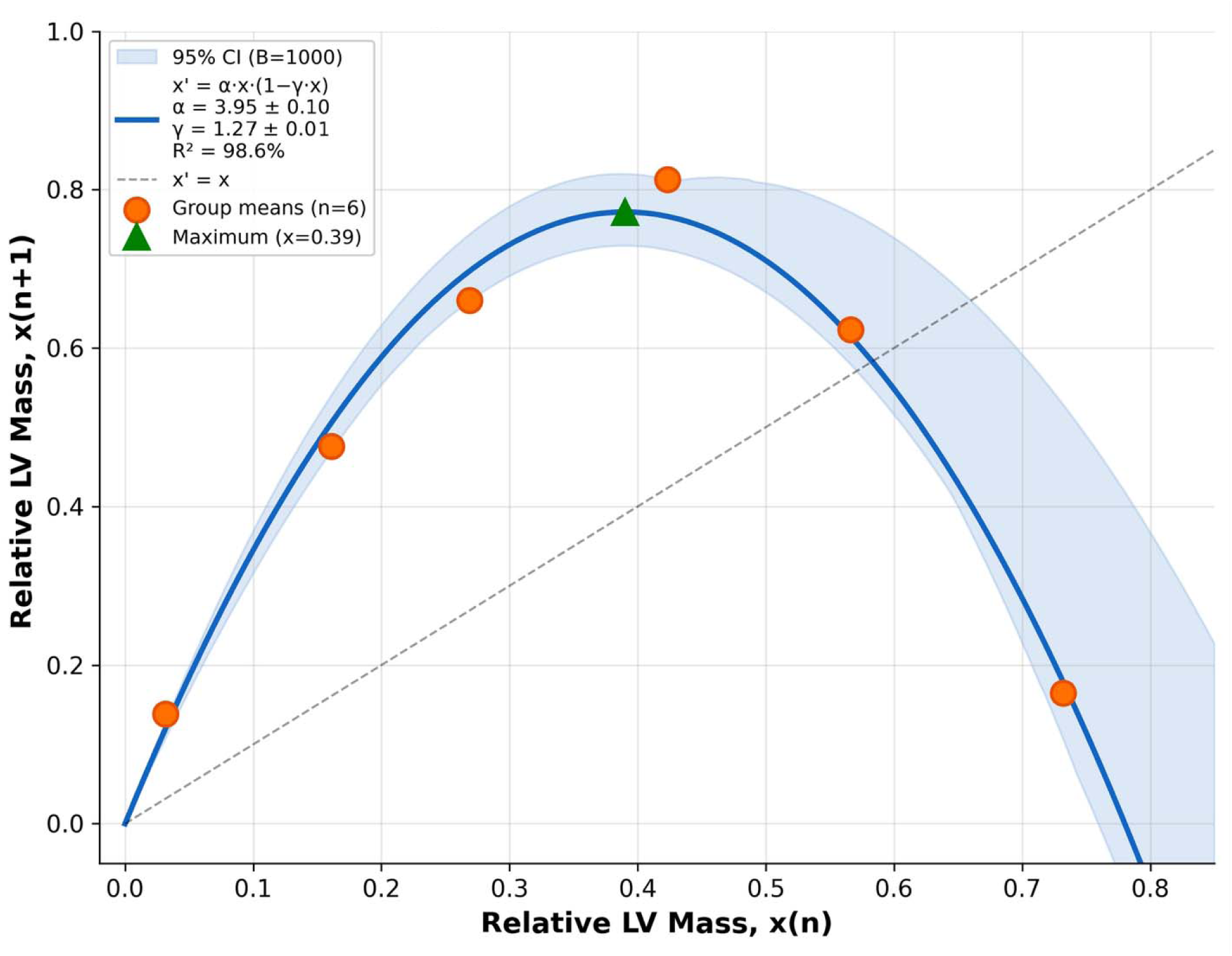
Logistic map of LV mass remodeling. Normalized LV mass at iteration n (x-axis) versus next-iteration mass at n+1 (y-axis) for *six* time-ordered group means. Solid curve: fitted logistic equation x′ = 3.952·x·(1−1.267·x). Dashed line: identity line (x′ = x). The intersection of the parabola with the identity line defines the fixed point x* = 0.59, corresponding to an LV mass of approximately 179 g (back-transformed via the inverse of the normalization x = LV mass/100 − 1.2). For α = 3.952, this fixed point is locally unstable. The value *of* α = 3.952 is compatible with a system operating near the threshold of complex nonlinear dynamics (α > 3.57), conditional on the assumption that the logistic equation is an adequate descriptor of the remodeling process; alternative concave models were not formally compared [14]. This is a population-level exploratory fit based on *six* grouped points rather than dense individual trajectories. The shaded blue area represents the bootstrap-generated confidence interval around the fitted line (nonparametric 1,000 resamples with replacement). The band is wider on the descending branch due to point distribution throughout the fitted line. **Alt text:** Scatter plot of six group-mean data points (labelled G-I to G-VI) fitted to an inverted parabolic logistic curve x′ = 3.952·x·(1−1.267·x). A diagonal dashed identity line intersects the parabola at the fixed point (x = 0.59). A light blue shaded band around the fitted curve represents the bootstrap 95 % confidence interval.

The fixed point x* = (α − 1) / (α · γ) = 0.59 corresponds to a back-transformed LV mass of approximately 179 g. For α = 3.952, |α · (1 − 2γx*)| = 1.95 > 1, indicating that the fixed point is locally unstable, as expected above the complexity threshold. The fixed point therefore serves as a mathematical reference separating two regions of the parabola, not as an attractor: patients with baseline x below x* operate on the ascending branch, those above on the descending branch.

### Lyapunov Exponent and Mortality

Using the 10-year endpoint, survivors exhibited positive LE (+0.349 ± 0.342, n = 40), indicating preserved dynamic complexity; nonsurvivors exhibited negative LE (−0.459 ± 0.377, n = 10; Mann-Whitney p < 0.001), indicating loss of complexity. The bootstrap distribution of ΔLE (LE_survivors − LE_nonsurvivors) (B = 1,000) was positive in 100 % of resamples (95 % CI [+0.59, +1.07]; Table 2A).

**Table 2.** Lyapunov exponent by outcome and multivariable regression of 10-year mortality.

| Panel A — Lyapunov exponent (LE) by outcome (N = 50) |  |  |  |  |  |  |  |  |
| --- | --- | --- | --- | --- | --- | --- | --- | --- |
|  | Survivors (n = 40) |  |  | Nonsurvivors (n = 10) |  |  | P* | Bootstrap ΔLE > 0 (B = 1,000) |
| LE (mean ± SD) | +0.349 ± 0.342 |  |  | −0.459 ± 0.377 |  |  | <0.001 | 100% |
| Panel B — Multivariable penalized regression of 10-year mortality (N = 50) |  |  |  |  |  |  |  |  |
|  | Logistic regression (Firth) |  |  | Cox regression (Firth, λ = 0.1) |  |  |  |  |
| Variable | β | OR [95% CI] | p | β | HR [95% CI] | p | PH p | Weight |
| EF < 51.7% | 4.5 | 88.3 [4.6–1690] | 0.003 | 2.3 | 10.14 [3.04–33.84] | 0.0002 | 0.696 | 2 |
| HRmax < 109 bpm | 3.3 | 25.8 [1.4–470] | 0.028 | 1.2 | 3.15 [1.05–9.47] | 0.041 | 0.334 | 1 |
| Model performance | AUC = 0.965 (apparent); optimism-corrected AUC = 0.959 [0.923–0.999] |  |  | C-index = 0.936 (apparent); optimism-corrected C-index = 0.929 [0.886–0.976] |  |  |  |  |

### Multivariate Regression and Risk Score

Among the 17 significant univariate predictors, collinearity analysis identified two groups: ventricular function (EF < 51.7 %, LVESD > 4.1 cm, LVEDD > 5.6 cm, LV mass > 217.9 g, Los Andes III; pairwise r = 0.63–0.89, forming a connected cluster above the |r| > 0.50 threshold) and ventricular arrhythmia (paired PVCs > 16/day, NSVT, PVCs > 655/day; r = 0.78−0.95). Time domain SAECG QRS abnormal and NYHA class II were excluded because all positive cases occurred exclusively among nonsurvivors. After retaining one representative per collinearity group (EF < 51.7 % and paired PVCs > 16/day), nine candidates were included in the Firth backward regression analysis. Seven variables were sequentially eliminated (SDNN < 41 ms, paired PVCs > 16/day, mitral regurgitation, SVE runs > 386/day, LA diameter > 4.4 cm, PSVT, and HR min > 71 bpm), yielding a final model with EF < 51.7 % (p = 0.003) and HRmax < 109 bpm (p = 0.028) (Table 2 Panel B). No eliminated variables regained significance after reintroduction.

#### Cox proportional hazards sensitivity analysis

The Cox-Firth model confirmed the logistic findings (Table 2B): EF < 51.7 % (HR = 10.14, 95 % CI 3.04 to 33.84, p < 0.001) and HRmax < 109 bpm (HR = 3.15, 95 % CI 1.05 to 9.47, p = 0.041), with the proportional-hazards assumption satisfied (Schoenfeld p = 0.696 and 0.334). The optimism-corrected C-index (0.929, 95% CI 0.886 to 0.976) was similar to the logistic AUC (0.959, 95% CI 0.923 to 0.999), and this convergence supports internal consistency rather than independent validation (see Limitations). Both models converged on identical 2:1 integer weights, yielding the same score: 2 × [EF < 51.7 %] + 1 × [HRmax < 109 bpm]. Bootstrap stability (B = 1,000) of Cox coefficients showed EF β > 0 in 100 % of resamples (p < 0.05 in 99.5 %) and HRmax β > 0 in 99.6 % of resamples (p < 0.05 in 51.8 %), consistent with the assigned weights.

### Clinical Risk Score and Lyapunov Exponent

Figure 2 shows the relationship between the clinical risk scores and Lyapunov exponents, comparing the study-derived score (Panels A and C) with the established Rassi score (Panels B and D). Both scoring systems showed monotonic mortality gradients, and LE declined monotonically across higher-risk strata.

**Figure 2 legend.**
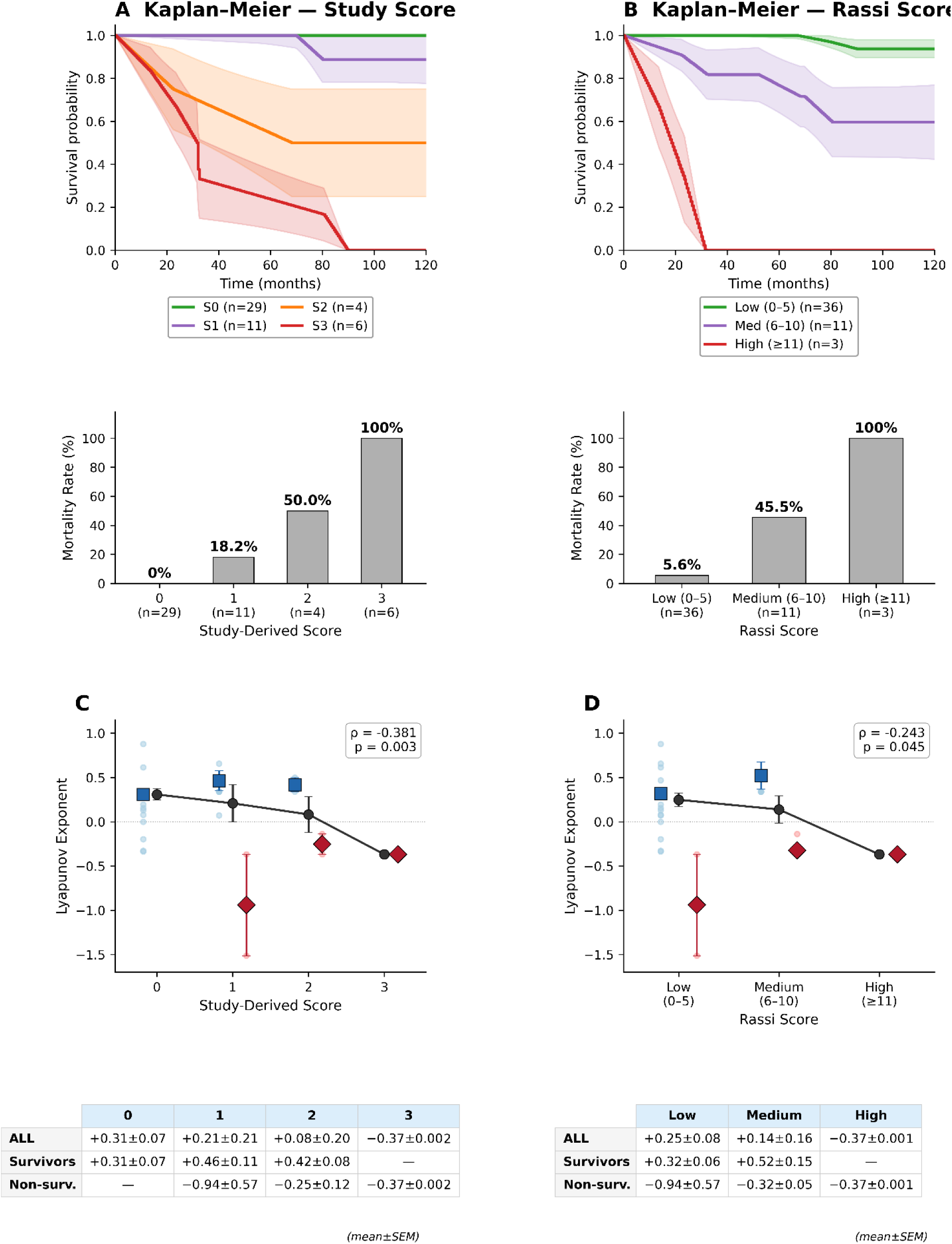
Clinical risk scores, Kaplan–Meier survival, mortality, and Lyapunov exponents. Panels A and B: Kaplan–Meier survival curves over the 120-month follow-up, stratified by the study-derived score (A; Table 2B) and the Rassi score (B). Color coding for Panels A and B: Score 0 green, Score 1 purple, Score 2 orange, Score 3 red; Rassi Low green, Medium purple, High red. Mortality rate bar plots (below Panels A and B, not re-lettered) display the 10-year cumulative mortality by stratum, with sample sizes annotated under each bar. The shaded bands represent the bootstrap standard error of the mean (B = 1,000 resamples with replacement, computed per stratum at each time point). Panels C and D: Mean Lyapunov exponent (± SEM) across strata of the study-derived score (C) and Rassi score (D). Black circles indicate all patients combined; blue squares indicate survivors (horizontally offset Δx = −0.18); red diamonds indicate nonsurvivors (offset Δx = +0.18). Pale blue and pale red points indicate individual patient LE values aligned with the respective group markers. Spearman correlation coefficients (one-sided, H : ρ < 0) are shown in the upper right corner of Panels C and D. Bottom tables (below Panels C and D, not re-lettered): LE values (mean ± SEM) per stratum for all patients, survivors, and nonsurvivors. In both scoring systems, survivors generally showed positive LE and nonsurvivors showed negative LE across risk strata. **Alt text:** Four-panel composite figure. Panels A and B display Kaplan–Meier survival curves over 120-month follow-up, stratified by the study-derived score (A) and the Rassi score (B), with mortality rate bar plots beneath. Panels C and D plot mean Lyapunov exponent (± SEM) across score strata for all patients (black circles), survivors (blue squares), and nonsurvivors (red diamonds), with one-sided Spearman correlation coefficients annotated. Survivors generally show positive Lyapunov exponents; nonsurvivors show negative values.

For the study-derived score (Score = 2 × [EF < 51.7 %] + 1 × [HRmax < 109 bpm], range 0–3), the mortality rate increased from 0 % (Score 0, n=29) to 18.2 % (Score 1, n=11), 50.0 % (Score 2, n=4), and 100 % (Score 3, n=6). The mean LE declined from +0.309 ± 0.351 (Score 0) to +0.209 ± 0.694 (Score 1), +0.083 ± 0.405 (Score 2), and −0.368 ± 0.002 (Score 3). Across the score strata, survivors generally showed positive LE, whereas nonsurvivors showed negative LE (Kruskal–Wallis H = 14.97, p = 0.002; Spearman ρ = −0.381, p = 0.003; bootstrap ρ<0 in 99.4 % of resamples).

The Rassi score [1] was computed for all 50 patients. The SAECG QRS amplitude was measured as the 3D vector magnitude (Vr = √3 × Vc), so the low-voltage threshold was applied as Vc < 0.866 mV (equivalent to Vr < 0.5 mV). Missing LVEDD values (n = 2) were imputed using the outcome group means (survivors 4.92 cm; nonsurvivors 5.98 cm).

The Rassi score replicated the pattern: positive LE in survivors, negative in nonsurvivors. Low-risk patients (score 0–5; n = 36, mortality 5.6 %) showed LE = +0.319 ± 0.333 in survivors vs. −0.940 ± 0.809 in nonsurvivors (p = 0.011); medium-risk (score 6–10; n = 11, mortality 45.5 %) showed +0.523 ± 0.375 vs. −0.322 ± 0.103 (p = 0.006); all three high-risk patients (score ≥ 11; mortality 100 %) had negative LE (−0.368 ± 0.001). Mortality increased and LE decreased monotonically across Rassi strata (overall LE +0.249 → +0.139 → −0.368; Kruskal-Wallis p = 0.020; Spearman ρ = −0.243, p = 0.045). Bootstrap validation (B = 1,000): Study Score ρ = −0.372 ± 0.150 (ρ < 0 in 99.4 % of resamples); Rassi ρ = −0.235 ± 0.158 (ρ < 0 in 92.4 % of resamples).

### Remodeling Trajectories

Among 35 patients with paired echocardiographic data (28 survivors, seven nonsurvivors at the 10-year endpoint; mean follow-up interval 5.9 ± 2.7 years), Figure 3 shows remodeling trajectories stratified by clinical outcome (group means ± SEM).

1. LV mass (Figure 3A). Nonsurvivors had higher LV mass than survivors at baseline (208.9 ± 59.9 vs. 142.3 ± 42.4 g, p = 0.006) and follow-up (259.7 ± 67.9 vs. 192.9 ± 69.1 g, p = 0.021); both groups showed progressive hypertrophy with persistent between-group separation.
2. EF (Figure 3B). Nonsurvivors had markedly lower EF at baseline (44.4 ± 17.7 vs. 74.2 ± 11.5 %, p = 0.001) and follow-up (35.0 ± 21.3 vs. 66.5 ± 13.0 %, p = 0.006); nonsurvivors began below the EF < 51.7 % threshold identified by the Firth model, survivors well above it.
3. LA diameter (Figure 3C). Nonsurvivors had larger LA diameter at baseline (4.17 ± 0.66 vs. 3.41 ± 0.54 cm, p = 0.014), with greater divergence at follow-up (4.68 ± 0.62 vs. 3.73 ± 0.44 cm, p = 0.001).

**Figure 3 legend.**
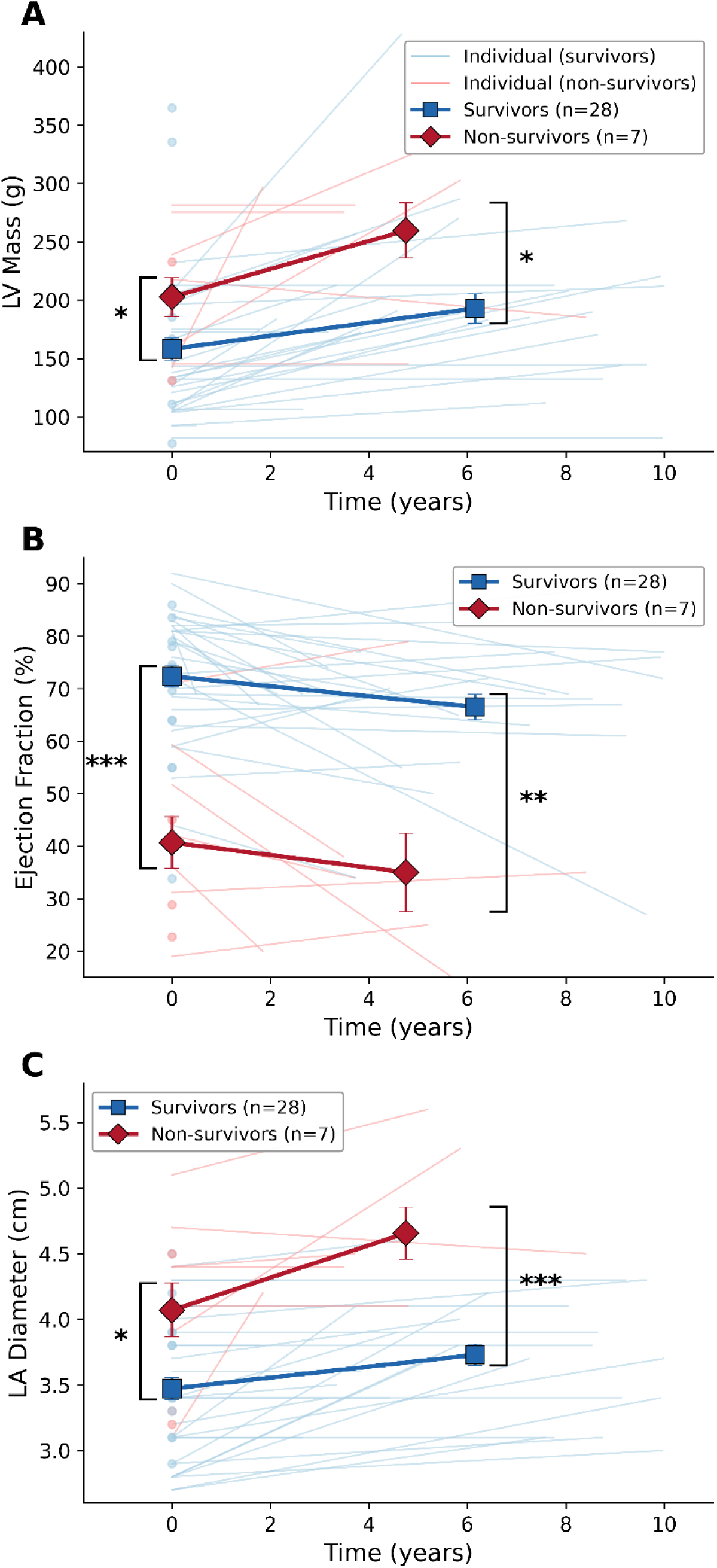
Remodeling trajectories based on clinical outcomes (10-year endpoint). Panel A: LV mass (survivors, n=28; nonsurvivors, n=7). Panel B: Ejection fraction (n=28 vs. n=7). Panel C: LA diameter (n=28 vs. n=7). Thin pastel lines represent individual patient trajectories connecting baseline and follow-up echocardiographic measurements; light blue, survivors; light red, nonsurvivors. Bold markers: group mean ± SEM. Blue squares: survivors. Red diamonds: Nonsurvivors. Brackets with significance symbols indicate Mann-Whitney comparisons at baseline (left) and follow-up (right): *p<0.05, **p<0.01. Panel A: LV mass — baseline 142.3 vs. 208.9 g (p=0.006), follow-up 192.9 vs. 259.7 g (p=0.021). Panel B: EF — baseline 74.2 % vs. 44.4 % (p=0.001), follow-up 66.5 % vs. 35.0 % (p=0.006). Panel C: LA — baseline 3.41 vs. 4.17 cm (p=0.014), follow-up 3.73 vs. 4.68 cm (p=0.001). The between-group differences were maintained from baseline to follow-up across all three parameters, consistent with the remodeling trajectories determined by the initial conditions. LA – left atrium. LV – left ventricle **Alt text:** Three-panel figure showing remodeling trajectories between baseline and 10-year follow-up. Panel A plots left ventricular mass (grams), Panel B plots ejection fraction (percent), and Panel C plots left atrial diameter (centimetres). In each panel, thin pastel lines connect baseline and follow-up values for individual patients (light blue: survivors; light red: nonsurvivors); bold blue squares and red diamonds mark group means ± SEM. Nonsurvivors show consistently higher LV mass and LA diameter and lower ejection fraction than survivors at both time points, with significance brackets marking Mann–Whitney comparisons.

Between-group separation was maintained from baseline to follow-up in all three parameters. The interpretation of this divergence — sensitive dependence on initial conditions vs. proportional progression from already divergent baselines — is addressed in the Discussion.

#### Variance analysis

LV mass variance increased from baseline to follow-up in both groups; EF and LA diameter variances were consistently greater in nonsurvivors. No Levene 4-group comparison reached significance (LV mass p = 0.625; EF p = 0.433; LA diameter p = 0.599), likely reflecting the small nonsurvivors sample size.

### Bootstrap Validation

Bootstrap distributions (B = 1,000) of the logistic map parameters (α, γ), equilibrium LV mass, and Lyapunov exponents according to survival status are presented in Supplementary Figure S1, acknowledging that group specific estimates rely on the limited number of nonsurvivors in the original sample. The bootstrap distribution of α was approximately normal (mean 3.96 ± 0.11, 95 % CI [3.76–4.17]), with 100 % of the resamples exceeding the complexity threshold (α > 3.57).

The bootstrap difference in LE between survivors and nonsurvivors (ΔLE = +0.80 ± 0.22) was positive in 100 % of the resamples, supporting the consistency of the complexity loss finding.

### Restricted Subset (Sensitivity Analysis)

When restricted to the subset in which both inter-patient derivatives were directly computed, the directional separation was preserved: survivors exhibited positive LE (+0.362 ± 0.552, n = 16) and nonsurvivors negative LE (−0.825 ± 0.972, n = 2; bootstrap ΔLE > 0 in 100 % of B = 1,000 resamples).

### Independent Validation in an Extended Follow-Up Sub-Cohort

Twelve patients of the cohort had additional echocardiograms obtained after the original 10-year endpoint, with a median extended observation of 17.6 years (range 10.4–19.8) beyond that endpoint. LV mass was computed by the Devereux 1986 formula at each serial measurement, and the intra-patient Lyapunov exponent was estimated from the longitudinal trajectory of each patient, both without and with normalization by the variable inter-examination intervals. The Spearman correlation between the intra-patient LE and the Rassi score was ρ = −0.66 (one-sided p = 0.010, n = 12; bootstrap ρ < 0 in 98.9 % of B = 1,000 resamples), recovering the same direction and magnitude as the inter-patient Rassi-LE gradient reported in the primary analysis (ρ = −0.243, one-sided p = 0.045, N = 50). Eleven of the 12 patients were 10-year survivors at the original endpoint; one died in 2020, beyond that endpoint. Per-patient values are reported in Supplementary Table S4.

## Discussion

This exploratory study provides preliminary evidence that long-term LV mass remodeling in chronic Chagas cardiomyopathy may follow a deterministic nonlinear pattern near the complex dynamical regime (α = 3.95) and that negative Lyapunov exponents may be associated with contemporaneous mortality. The convergence of two methodologically distinct frameworks (Firth-penalized logistic regression and Cox-Firth proportional hazards) on identical predictors, cutoffs, and 2:1 integer weights supports the score structure and reduces the likelihood of single-method artifact.

The retrospective design allowed patients to be grouped by their outcome, with each group sharing a common remodeling dynamic. Lyapunov exponents were computed from finite differences in LV mass between consecutive patients in admission order, separately at baseline and follow-up. This inter-group approach provides a local estimator of ensemble divergence and assumes within-group homogeneity of the remodeling dynamic. The validity of this inter-patient formulation rests on the ergodic-theoretic premise that, for a stationary dynamical system, the divergence rate measured between members of a population sampled at the same nominal time points is theoretically equivalent to the divergence rate that would be measured along a single patient’s trajectory if such longitudinal data were available [15].

To our knowledge, this is the first study to: 1) provide preliminary evidence that structural cardiac remodeling may follow the logistic equation, extending nonlinear dynamics theory from beat-to-beat HRV (millisecond timescale) to year-to-year ventricular mass dynamics; 2) compute Lyapunov exponents from echocardiographic trajectories rather than electrocardiographic time series; and 3) suggest that LE is associated with mortality in Chagas disease. Whether this loss of complexity reflects microscopic structural disorganization [16], such as progressive myofiber disarray from parasite persistence, chronic inflammation, microvascular injury, fibrosis, and autonomic dysregulation [1,2], remains to be tested by dedicated histological or imaging studies. Previous applications of nonlinear dynamics to Chagas disease focused on HRV entropy: Defeo et al. [10], Cornejo et al. [11], Valdez et al. [12], and the PhysioNet Challenge 2025 [17], all operating in the electrical-dynamics domain. The present study provides preliminary evidence that structural cardiac remodeling, a different biological process operating on a different timescale, may behave as a complex nonlinear dynamical system, extending this framework beyond the electrical domain [18,19,20].

### Lyapunov Exponent and the Complexity Loss Paradigm

Our findings extend the paradigm established by Poon and Merrill [5] and Wu et al. [7] from beat-to-beat HRV to annual structural remodeling. Both domains showed the same pattern: surviving hearts exhibited complex nonlinear dynamics (positive LE), whereas hearts of nonsurvivors lost complexity (negative LE). Poon and Barahona [8] showed that the noise limit, a quantity closely related to the Lyapunov exponent, can be titrated in cardiac time series, providing a framework for distinguishing deterministic complexity from stochastic fluctuations, consistent with our finding that the LE discriminates mortality. The related framework of “complexity variability” proposed by Valenza et al. [21], using the variance of instantaneous Lyapunov spectra, supports the broader paradigm that changes in cardiovascular complexity characterize disease, though the metric differs from the scalar LE used presently.

The 10-year endpoint provided the sharpest LE discrimination, although it was based on a small nonsurvivors sample because it captured deaths that occurred during the same period as the echocardiographic assessments. This temporal concordance positions LE as a contemporaneous marker of dynamic state, not a long-term predictor.

#### Biological Interpretation of the Logistic Parameters

The fitted parameters describe the population-level remodeling pattern and should not be mapped to specific biological processes. Any combination of pro-remodeling and counterregulatory mechanisms yielding the same macroscopic dynamics is expected to produce similar parameter values. The fitted α = 3.95 is compatible with a complex nonlinear regime. However, this compatibility should not be read as confirmation, given that the logistic equation was not formally compared with alternative concave models.

#### Empirical Convergence Consistent with the Theoretical Equilibrium

The mathematical equilibrium of the logistic equation under the fitted α and γ is x* = 0.59, corresponding to LV mass ≈ 179 g. For α = 3.952, this fixed point is locally unstable (|α · (1 − 2γx*)| = 1.95 > 1), as predicted for any α exceeding the complexity threshold of 3.57.

Population-level convergence is therefore not expected and was not observed: among 35 patients with paired echocardiograms, the mean absolute distance from the fixed point increased from 50.5 g at baseline to 55.7 g at follow-up (Wilcoxon signed-rank p = 0.95), and the six patients with baseline mass within 20 g of x* showed continued mass gain (+44 g on average) rather than stabilization. The outcome-stratified contrast was directionally informative (50 % of survivors moved closer vs. 14 % of nonsurvivors) but modest in magnitude. These observations are internally consistent with the dynamical regime: in the complex regime, trajectories diverge locally from the unstable fixed point and exhibit sensitive dependence on initial conditions rather than convergence.

#### Independent Validation by Intra-Patient Longitudinal Analysis

The extended sub-cohort provides an independent validation of the inter-patient formulation. The intra-patient Spearman correlation between LE and the Rassi score (ρ = −0.66, one-sided p = 0.010, n = 12; bootstrap ρ < 0 in 98.9% of B = 1,000 resamples) recovered the same direction and magnitude as the inter-patient Rassi-LE gradient reported in the primary analysis (ρ = −0.243, one-sided p = 0.045, N = 50). The two formulations, derived from independent analytical designs and from time scales an order of magnitude apart, recover the same Rassi-driven gradient in dynamic complexity, supporting the ergodic premise that motivated the inter-patient methodology.

#### Complementary Dimensions of Risk

The clinical risk score developed in this study and Lyapunov exponent capture complementary risk dimensions. The score combines ejection fraction < 51.7 % and maximum heart rate < 109 bpm on Holter monitoring, quantifying the severity of the baseline structural and autonomic substrates. In contrast, LE reflects the dynamic behavior of the remodeling process itself. LE declined monotonically across score strata in the cohort (Spearman ρ = −0.381, p = 0.003); the prespecified sensitivity check supported the same direction, reinforcing the robustness of the score-LE association.

This score was not intended to serve as an additional prediction tool for chronic Chagas cardiomyopathy. Established scores, including the Rassi score [1], SEARCH-Rio score [22] and SamiTrop risk model [23], were validated for clinical risk prediction in this population. The present score was developed to examine whether baseline characteristics, viewed as initial conditions of the proposed dynamical system, are linked to subsequent remodeling dynamics. Therefore, current findings are secondary and hypothesis-generating, complementing rather than competing with the established instruments.

The Rassi score replicated the directional pattern with non-overlapping predictors (Figure 2D): mortality rose monotonically across strata (5.6 % to 45.5 % to 100 %) and LE declined (+0.249 to +0.139 to −0.368; Spearman ρ = −0.243, p = 0.045). Within each stratum, survivors had positive LE and nonsurvivors negative LE. Concordance across two independently constructed scoring systems strengthens this association, pending confirmation in larger datasets.

Trajectory analysis (Figure 3) suggests that survivors and nonsurvivors differ primarily in initial conditions. Nonsurvivors started with higher LV mass (median 209 vs. 142 g, p = 0.006), lower EF (44 vs. 74 %, p = 0.001), and larger LA diameter (4.2 vs. 3.4 cm, p = 0.014), placing them on the descending branch of the fitted parabola versus the ascending-branch position of most survivors. These between-group separations were maintained at follow-up across all three echocardiographic parameters.

This observation can be interpreted within the framework of logistic dynamics, with caveats. In the equation x’ = α·x·(1−γ·x) with α = 3.952 and γ = 1.267, the parabola has a maximum at x = 1/(2γ), corresponding to LV mass ≈ 160 g under back-transformation, and the fixed point at LV mass ≈ 179 g is locally unstable. Baseline values below the parabolic maximum lie on the ascending branch of the parabola; values above it, including the fixed point, lie on the descending branch. Nonsurvivors began predominantly on the descending branch (median baseline 210 g), whereas survivors began predominantly on the ascending branch (median baseline 141 g). Notably, baseline LV mass alone was uncorrelated with empirical LE (Spearman ρ = −0.14, p = 0.34), and the LE separation between survivors and nonsurvivors (ΔLE = +0.81, p < 0.001) is not reducible to differences in LV mass alone. We emphasize that this interpretation is conditional on the logistic model being an adequate descriptor and cannot be distinguished, with the present design, from a simpler mechanism in which more severe disease at baseline progresses in proportion to its initial severity.

The increase in LV mass variance from baseline to follow-up (survivors 1,795 → 4,770; nonsurvivors 3,586 → 4,611) is directionally consistent with the expected behavior of a system near the threshold of complex nonlinear dynamics and extends the earlier observation of Benchimol-Barbosa et al. [24] of progressive intra-group dispersion. Given the small number of nonsurvivors with serial echocardiography, this pattern should be regarded as descriptive and hypothesis-generating.

#### Limitations

This study has some limitations. First, the 10-year follow-up included 10 nonsurvivors, constraining the precision of effect-size estimates; Post-Hoc Power Note (Supplementary Material) confirmed adequate power for both primary endpoints (LE d = 2.32 > minimum detectable 1.01; Spearman |ρ| = 0.381 > minimum detectable 0.35). Second, the inter-patient finite-difference Lyapunov formulation rests on within-outcome homogeneity of the remodeling dynamics, grounded in the ergodic premise (Methods); the prespecified sensitivity analysis (n = 18) preserved the directional separation. Third, the equilibrium LV mass (≈ 179 g) is a mathematical consequence of α and γ estimated from the same six grouped points and does not constitute an independent empirical validation of the logistic equation. Fourth, the large baseline differences between outcome groups (median EF 74 % vs. 44 %, LV mass 142 vs. 209 g) make the progressive between-group divergence compatible with proportional disease progression from divergent baselines, independently of any nonlinear mechanism. Fifth, the logistic equation was not formally compared with alternative concave models (e.g., Ricker map, Gompertz equation); its adequacy as a descriptor of LV mass dynamics remains to be tested. Sixth, the cohort was single-center and single-etiology; external validation in independent cohorts is required. Seventh, serial reclassification across staging systems [4,22] was not performed during follow-up; linking complexity trajectories to clinical progression remains to be addressed in future studies. Eighth, Pharmacological treatment during the 10 year follow up was at the physicians’ discretion and not systematically recorded, potentially influencing LV remodeling and outcomes. Finally, the Cox-Firth and Logistic Firth models share predictors and cutoffs; their convergence supports internal consistency rather than independent external validation.

### Conclusions

In this 10-year longitudinal cohort of patients with chronic Chagas cardiomyopathy, LV mass remodeling is compatible with a logistic population-level model, and a loss of dynamical complexity, captured by a negative Lyapunov exponent, is associated with mortality and with higher-risk strata of both a parsimonious study-derived score (EF < 51.7%, HRmax < 109 bpm) and the established Rassi score. The findings are consistent with, but do not establish, a nonlinear dynamic complex system remodeling regime. These results warrant replication in larger independent cohorts, direct comparison of the logistic equation against alternative concave models, and prospective validation of the clinical score as a descriptor of initial dynamical conditions. If confirmed, Lyapunov-based characterization of structural remodeling could provide a complementary dimension of risk stratification in Chagas cardiomyopathy beyond established scoring systems.

## Methods

### Study Design and Population

This substudy was a retrospective analysis of The Signal-averaged ElectrocArdiogram in Long Term Follow-up of Chronic CHagas Disease - RIO de Janeiro Cohort (SEARCH-Rio) cohort, comprising 50 consecutive outpatients with serologically confirmed Chagas disease from 1995 to 1999. Patients were classified according to the Los Andes system (I–III) [25]. This study was registered at ClinicalTrials.gov (registration ID: NCT01340963). The present retrospective analysis was approved by the Research Ethics Committee of Hospital Universitário Pedro Ernesto / Universidade do Estado do Rio de Janeiro (CEP-HUPE/UERJ) under protocol CAAE 97403926.4.0000.5259, with waiver of informed consent. The research was conducted in accordance with the Declaration of Helsinki.

### Clinical Follow-up

Information on clinical follow-up and subsidiary complementary evaluations has been described previously [22].

### Nonlinear Dynamics Analysis

The logistic-map fit was constructed at the population level by aggregating temporally proximate baseline LV mass measurements. Patients were sorted chronologically by date of admission, and the resulting sequence was partitioned into six contiguous, non-overlapping windows of variable width (windows were chosen so that each one contained at least 3 patients and that the range of admission dates within each window did not exceed 6 months) spanning the observation period; the choice of window boundaries balanced sample size within each window against the dispersion of admission dates. For each window, the mean baseline LV mass and the mean follow-up LV mass were computed, yielding six (mean-baseline, mean-follow-up) pairs. After normalization to relative LV mass [x = (LV mass / 100) − 1.2], these six time-ordered group means were fitted to the recursive logistic equation [13,14]: x’ = α·x·(1 − γ·x), where α is the logistic growth parameter and γ scales the normalized LV mass to the unit-parabola domain.

Values of α > 3.57 enter the complex dynamical regime through successive transitions to increasingly complex dynamics. This population-level fit was based on grouped time points and should be interpreted as an exploratory phenomenological model. The empirical Lyapunov exponent was computed as LE = log10(|derivative(n+1)| / |derivative(n)|) for each patient with serial LV mass measurements, with derivatives estimated as finite differences in LV mass between consecutive patients in admission order, separately at baseline and follow-up [26].

Because echocardiographic sampling was sparse (two time points per patient), this inter-patient finite-difference formulation was adopted in lieu of a full phase-space reconstruction; Under the assumptions of stationarity across the cohort and ergodicity of the underlying dynamical system, the inter-patient divergence rate measured here is theoretically equivalent to the intra-patient divergence that would be observed with multiple serial images per patient [15]. This ergodic premise is empirically supported by the extended follow-up sub-cohort analysis.

### Clinical Endpoints

The primary endpoint was all-cause mortality within 10 years of enrollment (40 survivors, 10 nonsurvivors; mean follow-up 96.3 ± 41.4 months, median 109.4 months), contemporaneous with the Lyapunov assessment period.

### Clinical Risk Scores

A clinical risk score for 10-year mortality was developed by Firth penalized logistic regression, which provides bias-reduced estimates for small samples with infrequent events. Candidate predictors included categorical variables significant by Fisher’s exact test (p < 0.05) and continuous variables significant by Mann-Whitney U (p < 0.05); the latter were dichotomized at optimal Youden cutoffs from receiver operating characteristic (ROC) curves. Collinearity was assessed by Pearson correlation prior to model entry (|r| > 0.50; lowest univariate p-value retained). Variables with univariate p < 0.10 entered the multivariate model. Backward elimination removed the variable with the highest p at each step until all predictors reached p < 0.05; eliminated variables were individually reintroduced to check for recovered significance.

Score weights were proportional to Firth coefficients, rounded to integer. The score was developed to explore the severity-transition behavior of the dynamical system rather than a clinical prediction tool.

### Cox proportional hazards sensitivity analysis

To address unequal follow-up times and provide time-aware effect estimates, a Cox proportional hazards model with Firth-type penalization (λ = 0.1, lifelines library) was fitted using the same two predictors retained by the logistic model. The proportional hazards assumption was assessed by Schoenfeld residuals with rank-transformed time. Model discrimination was quantified by Harrell’s concordance index (C-index); the apparent C-index was computed for the full sample, and the optimism-corrected C-index with 95 % CI was obtained by Harrell bootstrap with B = 1,000 resamples, matching the logistic-model AUC methodology. Cutoffs were not re-optimized to preserve methodological uniformity. Integer score weights followed a rounding rule (β to one decimal, then to integer).

Logistic regression remained the primary model given the temporal contemporaneity between the 10-year endpoint and the echocardiographic assessment period.

### Remodeling Trajectory Analysis

Remodeling trajectories were constructed for patients with paired echocardiographic measurements (LV mass, LVEF, LA diameter, n=35), stratified by clinical outcomes (survivors vs. nonsurvivors at the 10-year endpoint). The annual rate of change (slope) was computed as (follow-up value − baseline value) / interval for LV mass, ejection fraction, and left atrial diameter. These individual trajectories represent the fundamental parameters of the logistic dynamical system: LV mass captures the variable x in the logistic equation x’ = α·x·(1−γ·x), ejection fraction indexes the functional consequences of the remodeling process, and LA diameter reflects the hemodynamic burden of progressive ventricular dysfunction. Between-group differences in slopes and intercepts were assessed using the Mann-Whitney U test.

Variance heterogeneity across outcome groups and time points were evaluated by the Levene 4-group test (survivors and nonsurvivors at baseline and follow-up). The trajectories were also stratified by four equally spaced baseline LV mass groups (G I–IV) to assess whether initially homogeneous groups diverged over time. Variance amplification from baseline to follow-up was assessed using the Levene 8-group test (four mass groups × two time points). Within-group changes were evaluated using the Wilcoxon signed-rank test.

### Statistical Analysis

Categorical variables were compared by Fisher’s exact or chi-square test; continuous variables by Student’s t or Mann-Whitney U test, after normality assessment by Shapiro-Wilk. Holter parameters followed standard criteria [27]. Spearman’s rank correlation (ρ) assessed monotonic associations between clinical scores and LE; ρ was tested one-sided (H : ρ < 0). Kruskal-Wallis was used to assess differences in LE across score strata. Model discrimination was evaluated by AUC, Nagelkerke R², and F1-score. All analyses were validated by nonparametric bootstrap (B = 1,000), including Kaplan-Meier curves, the Firth logistic AUC, the Cox-Firth C-index, and Spearman’s ρ. A p < 0.05 was considered significant; tests were two-sided unless otherwise specified. When a patient’s follow-up LV mass was unavailable, the inter-patient derivative was estimated using the outcome-stratified group mean (survivors and nonsurvivors), preserving N = 50 in primary analyses. A prespecified sensitivity analysis repeated the same tests in the restricted subset (n = 18) in which both inter-patient derivatives were directly computed. An independent intra-patient validation was performed in 12 patients with extended follow-up beyond the 10-year endpoint (median 17.6 years, range 10.4 to 19.8); the Lyapunov exponent was estimated from the longitudinal trajectory of each patient, with and without normalization by the variable inter-examination intervals. Given the exploratory nature, no formal multiple-comparisons correction was applied; p-values are hypothesis-generating. Statistical code (Python 3.x) was developed with AI assistance (Claude, Anthropic) and verified by the authors.

### Supplementary Material

Supplementary materials available online comprise a consolidated Supplementary Material document (4 Supplementary Figures, bootstrap distributions, analytical flowchart, logistic-map fit quality, and empirical LE by outcome; Supplementary Table S1, empirical LE by outcome and Score-LE association; Supplementary Methods detailing analytical procedures and Supplementary Note on Post-hoc Power and Sensitivity Analyses, including minimum detectable effect sizes for the two endpoints). Supplementary Code (ten Python scripts, input dataset, and README) was developed with AI assistance (Claude, Anthropic) and verified by the authors.

## Acknowledgments

The authors thank Adriano de Moraes, MD, for performing electrocardiographic examinations at enrollment; Marcia B. Castier, MD, PhD, for performing echocardiographic examinations at enrollment and follow-up; and Francisco M. Albanesi Filho, MD (in memoriam), Marcelo I. Bittencourt, MD, PhD, and Silvia H.C. Boghossian, MD, PhD, for their role in the long-term clinical follow-up of participants.

Use of artificial intelligence. The authors used Claude (Anthropic) during the preparation of this manuscript to assist with language editing, formatting of figures and tables, and iterative refinement of statistical code (Python) for the bootstrap analyses and figure generation. All statistical computations, methodological decisions, interpretation of results, and final wording were reviewed, verified against source data, and approved by the authors, who take full responsibility for the content and integrity of the manuscript. No generative AI tool was used to produce original scientific content, generate references, or draft sections of the manuscript independently of author supervision.

## Author Contributions

P.R.B.-B. conceived and designed the study, performed data curation and formal statistical analysis, generated the figures, drafted the manuscript, and supervised the project. A.C.L.B.-B. contributed to data analysis, methodological refinement, and critical revision of the manuscript. C.G.C. contributed to methodology, software implementation of the analytical pipeline, and critical revision of the manuscript. B.K.K. provided study oversight, methodological guidance, and critical revision of the manuscript for important intellectual content. All authors reviewed and approved the final version of the manuscript.

## Data Availability Statement

All data and analysis scripts that support the findings of this study are provided as Supplementary Information. The de-identified consolidated dataset (Script_Input_Data_N50.xlsx) and ten Python analysis scripts with documentation (README) are included in the Supplementary Code package. The Python source code for logistic-map fitting, Lyapunov exponent computation, Firth penalized regression, Cox-Firth sensitivity analysis, and bootstrap validation is available with the Supplementary Material. Further information is available from the corresponding author upon reasonable request.

## Preprint Deposition

An earlier version of this manuscript has been posted at bioRxiv (DOI: 10.64898/2026.04.16.719099; posted April 20, 2026).

