## Supplementary figures, tables, methods for "Nonlinear Dynamics of Left Ventricular Mass Remodeling in Chagas Cardiomyopathy"

**Contents:**

Supplementary Figures S1–S4 (with legends)

Supplementary Table S1 (Lyapunov exponent by outcome and Score-LE association)

Supplementary Methods (detailed analytical procedures and formulas)

**Supplementary Figures and Tables**

| **Contents** | **Page** |
| --- | --- |
| Supplementary Figure S1 | 2 |
| Supplementary Figure S1 Legend | 3 |
| Supplementary Figure S2 | 4 |
| Supplementary Figure S2 Legend | 5 |
| Supplementary Figure S3 | 6 |
| Supplementary Figure S3 Legend | 7 |
| Supplementary Figure S4 | 8 |
| Supplementary Figure S4 Legend | 9 |
| Supplementary Table S1 | 10 |
| Supplementary Methods | 11 |
| Supplementary Table S2 | 25 |
| Supplementary Table S3 | 26 |
| Supplementary Table S4 | 31 |

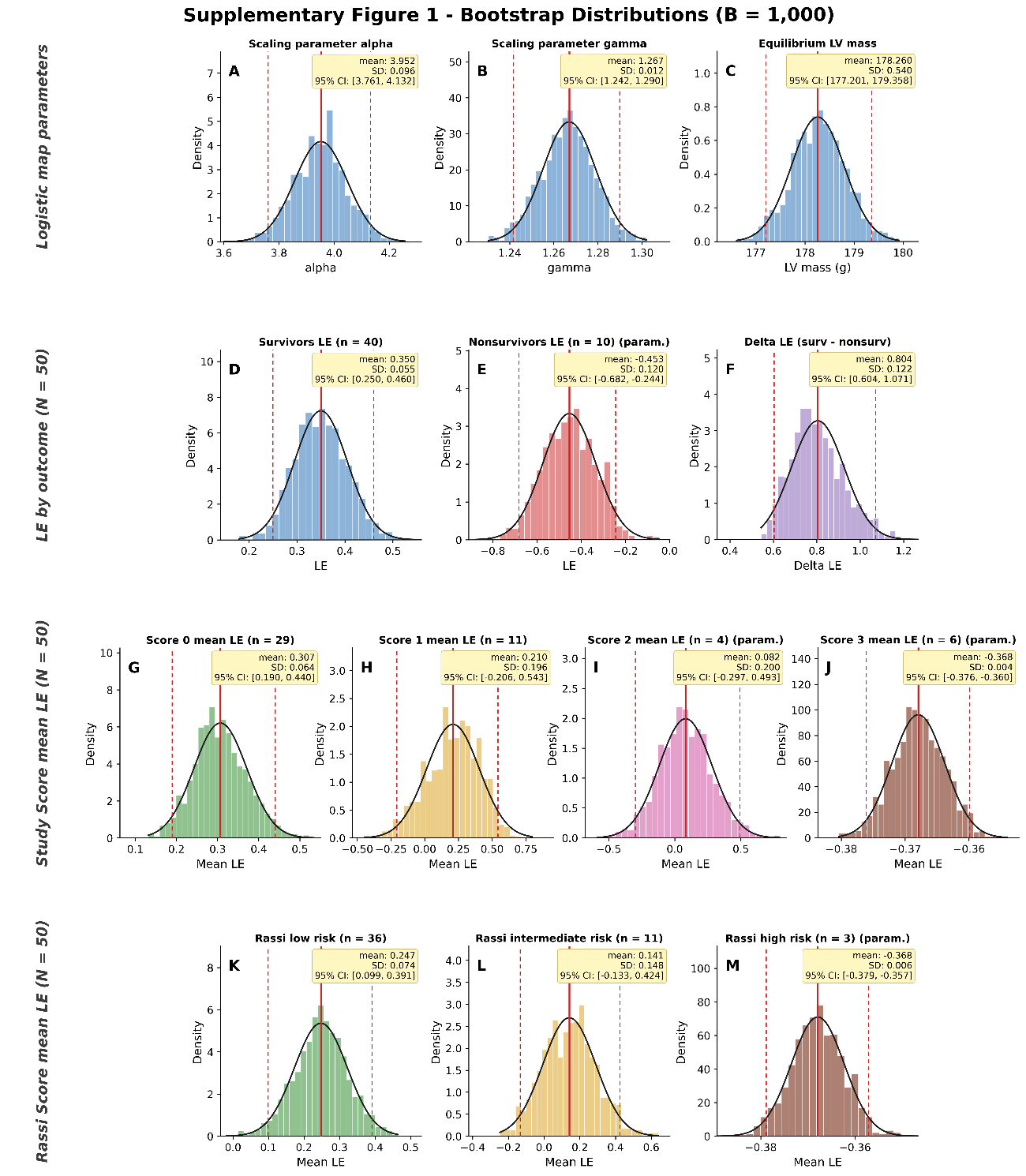

**Supplementary Figure S1**

**Supplementary Figure S1 Legend.** Bootstrap distributions (B = 1,000) for the cohort N = 50. Row 1, Panels A–C: logistic-map parameters obtained by residual (parametric) bootstrap from the 6-group nonlinear fit x′ = α·x·(1−γ·x), scaling parameter α (A), scaling parameter γ (B), and equilibrium LV mass x* = (α−1)/(α·γ) rescaled to grams (C). Row 2, Panels D–F: Lyapunov exponent by outcome, survivors (D, n = 40), nonsurvivors (E, n = 10), and ΔLE (F; ΔLE > 0 in 100 % of resamples). Row 3, Panels G–J: mean LE by Study-Derived Score, Score 0 (G, n = 29), Score 1 (H, n = 11), Score 2 (I, n = 4), and Score 3 (J, n = 6). Row 4, Panels K–M: mean LE by Rassi class, low (K, n = 36), intermediate (L, n = 11), and high (M, n = 3). Black curves: fitted Gaussian densities. Red solid and dashed lines: bootstrap mean and 95 % percentile confidence interval. Yellow boxes report the bootstrap mean, the bootstrap standard deviation (SD), and the 95 % CI. Panels marked “param.” (E, I, J, M) use parametric residual gaussian bootstrap when nonparametric resampling is unstable. The x-axis range of every panel was set to the bootstrap mean ± 3.8·SD.

***Alt text:*** *Composite multi-panel figure (A*–*M) showing bootstrap distributions of logistic-map parameters, Lyapunov exponents by outcome, and Lyapunov exponents by score stratum. Each panel is a histogram with an overlaid fitted Gaussian density curve, bootstrap mean and 95% confidence interval lines, and annotated summary statistics.*

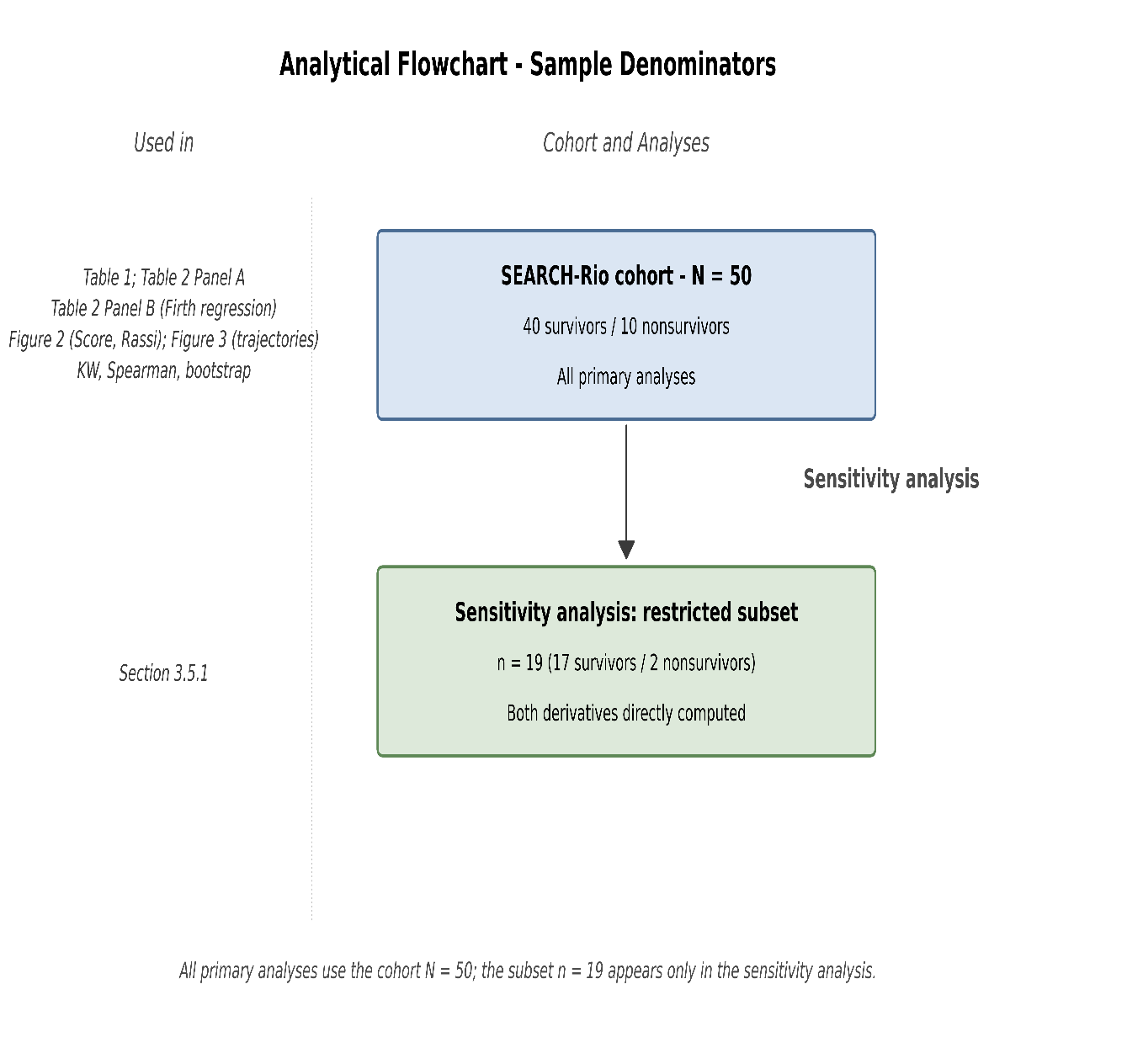

**Supplementary Figure S2**

**Supplementary Figure S2 Legend.** Analytical flowchart of the cohort and analyses. Data from the SEARCH-Rio cohort (N = 50; 40 survivors, 10 nonsurvivors) supports all primary analyses, including Tables 1 and 2 (Panels A and B), Figure 2 (Score and Rassi), Figure 3 (remodeling trajectories), and the Kruskal-Wallis, Spearman, and bootstrap tests. A prespecified sensitivity analysis examines robustness using the restricted subset (n = 18; 16 survivors, 2 nonsurvivors) for which both inter-patient derivatives were directly computed (Section 3.5.1).

***Alt text:*** *Flowchart with one upper box (SEARCH-Rio cohort, N = 50, used in all primary analyses) and one lower box connected by an arrow (restricted subset, n = 18), labelled as the prespecified sensitivity analysis.*

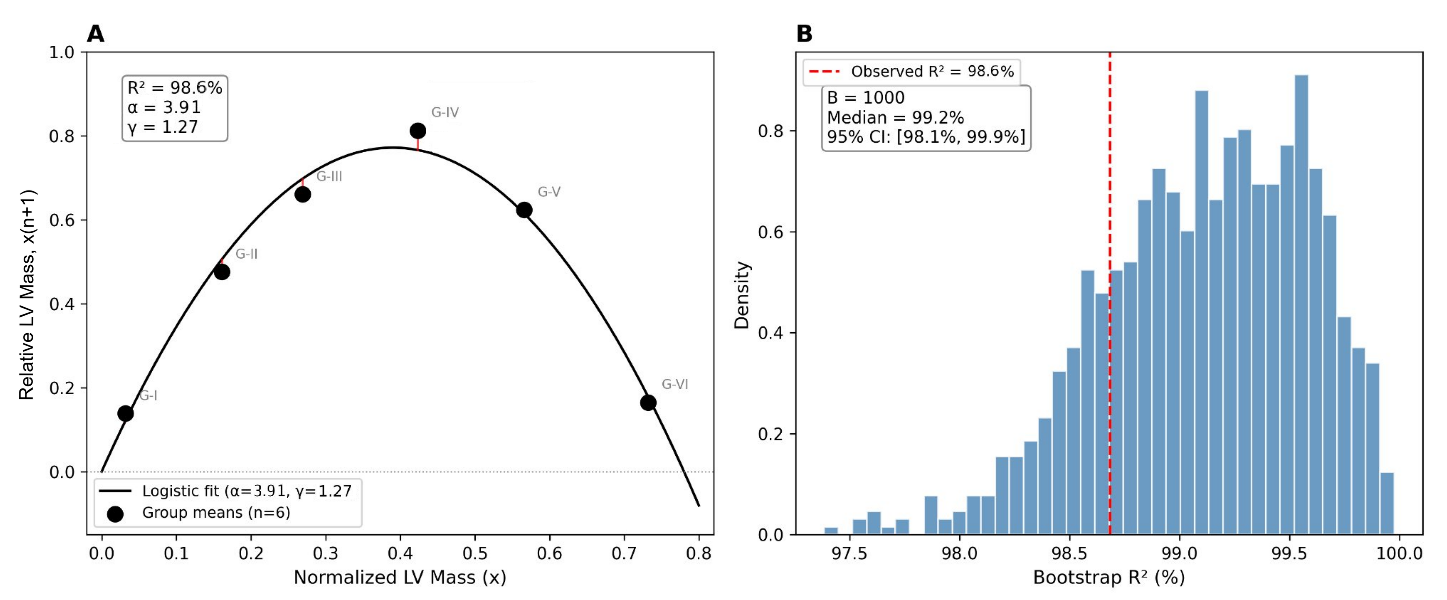

**Supplementary Figure S3**

**Supplementary Figure S3 Legend.** Logistic-map fit quality assessment. Panel A: measured group means (n = 6 time-ordered points) versus fitted logistic curve x′ = 3.952·x·(1−1.267·x), showing the 6 data points and fitted parabola with residuals. Panel B: bootstrap distribution of R² values (B = 1,000 residual resamples) demonstrating the stability of the model fit despite the limited number of fitted points.

***Alt text:*** *Two-panel figure. Left: six data points (G-I to G-VI) fitted to a parabolic logistic curve with short red residual segments connecting points to the curve. Right: histogram of 1,000 bootstrap R² values ranging from approximately 97.5% to 100%, with a vertical red dashed line marking the observed R² = 98.6%.*

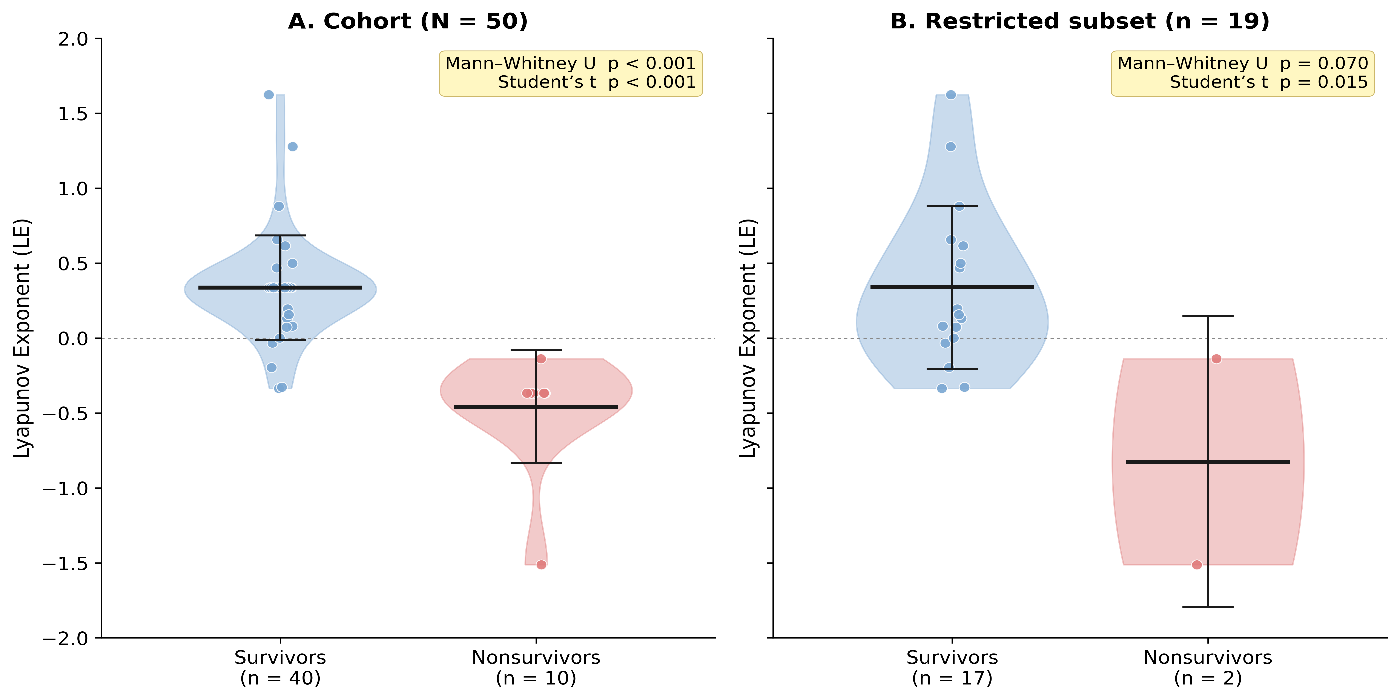

**Supplementary Figure S4**

**Supplementary Figure S4 Legend.** Empirical Lyapunov exponent by outcome. Panel A: cohort N = 50 (primary). Panel B: restricted subset n = 18 (sensitivity). Violin plots show the kernel density estimate of LE; thick horizontal bars indicate the mean and the vertical bars ±1 SD; individual data points (jittered) are overlaid. Survivors in blue, nonsurvivors in red. Statistical test p-values are shown in the upper-right corner of each panel.

***Alt text:*** *Two side-by-side violin plots comparing Lyapunov exponents between survivors (blue) and nonsurvivors (red). Panel A shows the cohort N = 50; Panel B shows the restricted subset n = 18. Survivors have positive LE, nonsurvivors negative.*

| **Analysis** | **n (surv / nonsurv)** | **LE survivors (mean ± SD)** | **LE nonsurvivors (mean ± SD)** | **MW U (primary)** | **Student’s *t* (secondary)** | **Spearman ρ Score-LE (*p*)** | **Bootstrap ΔLE > 0** |
| --- | --- | --- | --- | --- | --- | --- | --- |
| Primary (N = 50) | 40 / 10 | +0.349 ± 0.342 | −0.459 ± 0.377 | < 0.001 | < 0.001 | −0.381 (0.003) | 100 % |
| Sensitivity (n = 18) | 16 / 2 | +0.362 ± 0.552 | −0.825 ± 0.972 | 0.078** | 0.016 | +0.058 (0.818) | 100 % |

**Supplementary Table S1. Empirical Lyapunov exponent by outcome and Score-LE association: cohort N = 50 (primary) and restricted subset n = 18 (sensitivity).**

Empirical Lyapunov exponent (LE) by outcome and association with the study-derived score, in the cohort N = 50 (primary) and the restricted subset n = 18 (sensitivity). The directional separation between survivors and nonsurvivors and the bootstrap ΔLE > 0 in 100 % of resamples are preserved in both. Bootstrap: B = 1,000 resamples with replacement.** Student’s *t* teste p = 0.015

**SUPPLEMENTARY METHODS**

*Nonlinear Dynamics of Left Ventricular Mass Remodeling
in Chagas Cardiomyopathy*

Detailed Formulas and Computational Procedures

| **Contents** | **Page** | |
| --- | --- | --- |
| 1. Logistic Map of LV Mass Remodeling | 13 | |
| 1.1 Data preparation and normalization | 13 | |
| 1.2 Group mean computation (6 time-ordered points) | 13 | |
| 1.3 Nonlinear least-squares fitting | 13 | |
| 1.4 Fixed-point equilibrium | 14 | |
| 1.5 Bootstrap validation (residual method) | 14 | |
| 2. Empirical Lyapunov Exponent | 16 | |
| 2.1 Inter-patient divergence method | 16 | |
| 2.2 Derivatives (Deriv0, Deriv1) | 16 | |
| 2.3 LE computation (log₁₀) | 16 | |
| 2.4 Derivative estimation rule | 18 | |
| 2.5 Sensitivity analysis | 18 | |
| 3. Firth Penalized Logistic Regression | 19 | |
| 3.1 Univariate screening | 19 | |
| 3.2 Collinearity resolution | 19 | |
| 3.3 Backward stepwise elimination | 19 | |
| 3.4 Final model and clinical risk score | 20 | |
| 3.5 Score performance metrics | 20 | |
| 4. Cutoff Selection and Score Weight Derivation | 21 | |
| 4.1 Youden-Index Optimization of Continuous Cutoffs | 21 | |
| 4.2 Two-Step Half-Up Rounding Rule for Score Weights | 21 | |
| 5. Cox-Firth Proportional Hazards Sensitivity Analysis | 22 | |
| 5.1 Rationale | 22 | |
| 5.2 Model Specification | 22 | |
| 5.3 Proportional Hazards Assumption | 22 | |
| 5.4 Discrimination and Optimism Correction | 22 | |
| 5.5 Coefficient Stability | 22 | |
| 5.6 Convergence of Modeling Frameworks | 22 | |
| 6. Logistic Firth Bootstrap Optimism Correction | 23 | |
| 7. Post-hoc Power and Sensitivity Analyses | 24 | |
| 7.1 Background and rationale | 24 | |
| 7.2 Methods | 24 | |
| 7.3 Results | 26 | |
| 7.4 Interpretation | 28 | |
| 7.5 References | 30 | |
| 8. Independent Validation in an Extended Follow-Up Sub-Cohort | | 30 |
| 8.1 Rationale | | 30 |
| 8.2 Methods | | 30 |
| 8.3 Results | | 30 |
| 8.4 Significance | | 30 |

### 1. Logistic Map of LV Mass Remodeling

#### 1.1 Data Preparation and Normalization

The logistic map analysis uses paired echocardiographic LV mass measurements: baseline (LV_mass (LV mass at baseline), at admission) and follow-up (LV_mass_FU (LV mass at follow-up), at end of follow-up or last exam before death). The dataset comprises N=50 patients from the SEARCH-Rio cohort, of whom 35 have real serial echocardiographic data and 15 have follow-up values estimated by outcome-stratified group means.

#### Normalization procedure:

#### Each LV mass value (in grams) is normalized to a dimensionless relative mass by:

#### xᵢ = LV_massᵢ/100 − 1.2

#### x′ᵢ = LV_mass_FUᵢ/100 − 1.2

#### The offset 1.2 is a convenient arbitrary constant that places the values observed in this cohort approximately on the interval where the logistic equation x′ = α·x·(1 − γ·x) is conventionally defined. The same normalization is applied uniformly to baseline and follow-up values and is also used (inverted) to back-transform the fixed-point equilibrium to original mass units in Section 1.4, by LV_mass = (x + 1.2) · 100.

#### 1.2 Group Mean Computation (6 Time-Ordered Points)

The logistic map is constructed at the population level (inter-patient), not from individual patient trajectories. The procedure is as follows:

Step 1: Sort patients chronologically by date of admission within the observation window.

Step 2: Partition the sorted sequence into six contiguous, non-overlapping windows of variable width spanning the observation period; window boundaries balance sample size within each window against the dispersion of admission dates. The empirical partition adopted in the source dataset has varying window sizes across all analyzed patients with temporally adjacent computational columns available.

Step 3: For each group G_k, compute the mean of the normalized baseline values and the mean of the normalized follow-up values:

x̄(n)_k = mean{ xᵢ : i ∈ G_k }

x̄(n+1)_k = mean{ x′ᵢ : i ∈ G_k }

This yields 6 coordinate pairs (x̄(n)_k, x̄(n+1)_k), k = 1,...,6, which trace the iterative map. Low-mass groups (G1–G4) typically show x̄(n+1) > x̄(n) (ascending branch of the parabola), while high-mass groups (G5–G6) may show x̄(n+1) < x̄(n) (descending branch), which is the signature of the nonlinear regime.

#### 1.3 Nonlinear Least-Squares Fitting

The generalized logistic equation with two parameters is:

**x′ = α · x · (1 − γ · x)**

where α is the growth-rate parameter (bifurcation parameter) and γ is a scaling coefficient. When γ = 1, this reduces to the standard logistic map x′ = α · x · (1 − x).

The parameters α and γ are estimated by nonlinear least-squares minimization applied to the 6 group means:

minimize ∑ₖ₌₁⁶ [ x̄(n+1)_k − α · x̄(n)_k · (1 − γ · x̄(n)_k) ]²

Implementation: scipy.optimize.least_squares with bounds α ∈ [0, 10] and γ ∈ [0, 10], initial guess α₀ = 3.5, γ₀ = 1.2.

Goodness of fit:

R² = (Pearson r)², where r is the Pearson’s correlation coefficient

Results: α = 3.952 ± 0.096, γ = 1.267 ± 0.012, R² = 98.6%.

*The dynamical regime is determined by α: α < 3.0 → convergence to fixed point; 3.0 < α < 3.57 → periodic oscillations (bifurcation cascade); α ≥ 3.57 → regime of complex nonlinear dynamics (sensitive dependence on initial conditions). The finding α = 3.952 places the system in the regime of complex nonlinear dynamics*

#### 1.4 Fixed-Point Equilibrium

#### The fixed point x* is the value where x′ = x:

## α · x* · (1 − γ · x*) = x*

## x* = (α − 1) / (α · γ)

#### For α = 3.952, γ = 1.267: x* = (3.952 − 1) / (3.952 × 1.267) = 0.59.

#### Back-transforming to original mass units via the inverse of the normalization defined in Section 1.1:

#### LV mass* = (x* + 1.2) · 100 ≈ 179 g

#### Local stability of the fixed point is determined by the magnitude of the derivative of the map at x*: |α · (1 − 2γx*)| = |α − 2| = 1.95. Since this exceeds 1, the fixed point is locally unstable, consistent with α exceeding the complexity threshold of 3.57. In this regime, trajectories diverge locally from the fixed point and exhibit sensitive dependence on initial conditions; the fixed point therefore serves as a mathematical reference, not as an attractor.

#### 1.5 Bootstrap Validation (Residual Method)

The bootstrap uses the residual resampling method (Method B), which keeps the 6 x-values fixed and resamples only the residuals. This produces symmetric (bell-shaped) distributions for all parameters.

Procedure (B = 1,000 resamples):

Step 1: Compute residuals from the original fit: e_k = x̄(n+1)_k − α̂ · x̄(n)_k · (1 − γ̂ · x̄(n)_k), k=1,...,6.

Step 2: For each bootstrap iteration b = 1,...,1000: (a) Draw 6 residuals with replacement from {e₁,...,e₆}. (b) Construct pseudo-observations: y*_k = ŷ_k + e*_k, where ŷ_k is the fitted value. (c) Re-fit the logistic equation to (x̄(n)_k, y*_k), obtaining α*(b), γ*(b).

Step 3: Compute x**(b) = (α*(b) − 1) / (α*(b) · γ*(b)) for each resample.

Results:

| Parameter | Mean ± SD | 95% CI | Skewness | % > threshold |
| --- | --- | --- | --- | --- |
| α | 3.946 ± 0.102 | [3.76, 4.17] | 0.097 | 100% > 3.57 |
| γ | 1.281 ± 0.014 | [1.26, 1.30] | −0.363 | — |
| x* | 0.583 ± 0.005 | [0.57, 0.59] | 0.113 | — |

*The bootstrap mean α = 3.946 differs from the point estimate α = 3.952 by 0.15 %, within the expected variability for n = 6 data points and consistent with the symmetric residual bootstrap procedure described in this section.*

### 2. Empirical Lyapunov Exponent

#### 2.1 Inter-Patient Divergence Method

The empirical Lyapunov exponent (LE) quantifies the rate of divergence (or convergence) of neighboring trajectories in the remodeling space. Unlike the analytical LE derived from the logistic map derivative [LE = ln|α·(1−2γ·x)|], the empirical LE is computed from the actual patient data using an inter-patient divergence approach.

The LE is computed from the consolidated dataset Script_Input_Data_N50.xlsx (sheet 'Data_N50'), which contains all 50 patients of the SEARCH-Rio cohort with baseline (LV_mass) and follow-up (LV_mass_FU) values. Patients are processed in the analytic ordering by admission date. The restricted subset M18 (n = 18, 16 survivors / 2 non-survivors) consists of patients for whom both the patient and the immediate chronological predecessor have measured baseline AND follow-up masses, yielding directly computable Deriv0 and Deriv1. The remaining 32 patients have Deriv0 and Deriv1 estimated from outcome-stratified M18 group means at the derivative level (see Section 2.4), preserving the cohort size of N = 50 in primary analyses.

#### 2.2 Derivatives (Deriv0, Deriv1)

Patients are sorted chronologically by admission date. For consecutive patients i and i−1:

**Deriv0ᵢ = | LV_mass(i) − LV_mass(i−1) |**

**Deriv1ᵢ = | LV_mass_FU(i) − LV_mass_FU(i−1) |**

Deriv0 measures the inter-patient difference in baseline LV mass between consecutive patients (ordered by admission). Deriv1 measures the inter-patient difference in follow-up LV mass between those same consecutive patients. If the system is sensitive to initial conditions, small Deriv0 values can produce large Deriv1 values (divergence), and vice versa.

***Key distinction:*** *The logistic map is constructed inter-patient (group means). The LE is also inter-patient (consecutive-patient divergence), NOT intra-patient (within a single patient's trajectory over time). Under the assumptions of (i) stationarity of the chronic Chagas remodeling process across the cohort during the observation window and (ii) ergodicity of the underlying dynamical system, the inter-patient divergence rate measured here is theoretically equivalent to the intra-patient divergence that would be observed if each patient had been serially imaged at multiple time points (cf. Wolf et al., Physica D 1985; Eckmann & Ruelle, Rev Mod Phys 1985). The inter-patient formulation exploits the population as a delay-embedded reconstruction of the dynamical attractor.*

#### 2.3 LE Computation (log₁₀)

The empirical LE for each patient i (i > 1) is:

**LEᵢ = log₁₀( Deriv1ᵢ / Deriv0ᵢ )**

Conditions:

• Both Deriv0ᵢ > 0 and Deriv1ᵢ > 0 are required. If either is zero, LE is undefined for that patient.

• The first patient in the admission-ordered sequence has no LE (no predecessor).

Interpretation:

• LE > 0: Divergence — the difference in follow-up masses is greater than the difference in baseline masses; small differences at baseline amplify over time (nonlinear complex signature).

• LE < 0: Convergence — the difference in follow-up masses is smaller than at baseline; trajectories converge (loss of dynamical complexity).

• LE = 0: Neither divergence nor convergence.

*CRITICAL: The empirical LE uses log₁₀ (base 10), NOT ln (natural logarithm). This was verified by matching all 19 measured LE values in the source spreadsheet: log₁₀ matched 19/19 patients; ln matched 0/19.*

#### 2.4 Derivative Estimation Rule

When a patient’s follow-up LV mass was not available, the corresponding inter-patient derivative was estimated using the outcome-stratified group mean (survivors and nonsurvivors), preserving the cohort size of N = 50 in primary analyses. Estimation was performed at the derivative level, not at the LE level. The 18 patients of the restricted subset M18 retained their directly computed Deriv0 and Deriv1 values; the remaining 32 patients received the outcome-stratified M18 group means as Deriv0 and Deriv1, yielding a constant LE per outcome group for these patients (LE = +0.341 in survivors, LE = −0.368 in nonsurvivors).

#### 2.5 Sensitivity Analysis

A prespecified sensitivity analysis repeated the score-LE associations (KW, Spearman, bootstrap) in the restricted subset (n = 18) for which both inter-patient derivatives were directly computed. Results are reported in Supplementary Table S1.

### 3. Firth Penalized Logistic Regression

Endpoint: 10-year all-cause mortality (10/50 events).

All echocardiographic variables were dichotomized at their optimal Youden-index cutoff derived from individual ROC curves against the mortality endpoint:

| Variable | Cutoff | Direction | Univariate p |
| --- | --- | --- | --- |
| EF | 51.7% | < | <0.001 |
| LVESD | 4.1 cm | > | <0.01 |
| LVEDD | 5.6 cm | > | <0.01 |
| HR max | 109 bpm | < | 0.064 (MW) |
| LA diameter | 4.4 cm | > | <0.05 |
| SDNN | 41 ms | < | <0.05 |

Only variables with univariate p < 0.15 entered the multivariate model.

#### 3.2 Collinearity Resolution

Collinearity was assessed by Spearman correlation. Variables with |ρ| > 0.50 were grouped, and only the variable with the lowest univariate p-value was retained from each group:

Group 1 (ventricular function): EF < 51.7%, LVESD > 4.1 cm, LVEDD > 5.6 cm, LV_mass > 217.9 g (ρ = 0.50–0.87). Retained: EF < 51.7% (lowest p).

Group 2 (ventricular arrhythmia): paired PVCs > 16/24h, NSVT episodes ≥ 1 (NSVT01), PVC count > 655/24h (ρ = 0.78–0.95). Retained: paired PVCs > 16/24h (lowest p).

Independent variables: NYHA class ≥ II, paroxysmal SVT episodes, mitral regurgitation, LA diameter > 4.4 cm, supraventricular ectopy > 386/24h, HRmax < 109 bpm, SDNN < 41 ms.

#### 3.3 Backward Stepwise Elimination

Firth penalized logistic regression corrects the small-sample bias of maximum likelihood estimation by adding a Jeffreys prior penalty to the likelihood function:

L*( β) = L(β) · |I(β)|^1/2^

where L(β) is the standard likelihood and I(β) is the Fisher information matrix. This prevents infinite coefficient estimates in cases of quasi-complete separation (common with few events and multiple predictors).

Backward elimination procedure (fixed default):

Step 0: Enter all candidates after collinearity removal (7–9 variables). Step k: Fit Firth regression with current variables. Identify the variable with the highest p-value. If p > 0.05, remove that variable and repeat. Continue until all remaining variables have p < 0.05.

Elimination sequence: paired PVCs > 16/24h → supraventricular ectopy > 386/24h → paroxysmal SVT episodes → LA diameter > 4.4 cm → mitral regurgitation. No eliminated variable recovered significance upon reintroduction.

#### 3.4 Final Model and Clinical Risk Score

The final Firth regression model retained two variables:

| Variable | β (Firth) | SE | OR | 95% CI | p |
| --- | --- | --- | --- | --- | --- |
| EF < 51.7% | 4.5 | 1.545 | 88.3 | 4.6–1690 | 0.003 |
| HR max < 109 bpm | 3.3 | 1.519 | 25.8 | 1.4-470 | 0.028 |

The clinical risk score is derived from the regression weights:

**Score = 2 × (EF < 51.7%) + 1 × (HRmax < 109 bpm)**

Integer weights were derived by a two-step half-up rounding rule applied uniformly to both predictors (see Section 4.2): EF β = +4.481 → 4.5 → 5; HRmax β = +3.250 → 3.3 → 3; ratio 5:3 collapses to final integer weights 2:1. An independent Cox-Firth sensitivity model (Section 5) yielded β = +2.317 and +1.148 for EF and HRmax respectively, which under the same rounding rule produce identical 2:1 weights, confirming score robustness across modeling frameworks. Score range: 0–3.

#### 3.5 Score Performance Metrics

Model-level performance:

AUC apparent = 0.965; bootstrap optimism-corrected AUC = 0.959, 95% CI [0.923, 1.007] (B = 1,000 resamples).

Score ≥ 2 (optimal threshold):

Sensitivity = 0.800, Specificity = 0.950, PPV = 0.800, NPV = 0.950, F1 = 0.800.

Mortality gradient by score level:

| Score | n | Mortality (%) | LE (mean) |
| --- | --- | --- | --- |
| 0 | 29 | 0% | +0.309 |
| 1 | 11 | 18.2% | +0.209 |
| 2 | 4 | 50% | +0.083 |
| 3 | 6 | 100% | −0.368 |

The monotonic mortality gradient (0% → 18.2% → 50% → 100%) with parallel decline in LE validates the Score as a bridge between clinical risk stratification and nonlinear dynamics. Kruskal-Wallis (Score vs LE): H = 14.97, p = 0.002. Spearman ρ (Score vs LE) = −0.381, p (one-sided) = 0.003.

Concordance with the Rassi score was independently confirmed: Kruskal-Wallis p = 0.020 (Low/Medium/High: 5.6% → 45.5% → 100% mortality; LE: +0.249→ +0.139 → −0.368).

### 4. Cutoff Selection and Score Weight Derivation

#### 4.1 Youden-Index Optimization of Continuous Cutoffs

Each continuous candidate predictor entering the multivariate Firth model was dichotomized at its Youden-optimal cutoff, determined from its univariate ROC curve against the 10-year mortality endpoint (10/50 events). The Youden index J was defined as:

J(c) = Sensitivity(c) + Specificity(c) − 1

and maximized by exhaustive search across all observed values of the predictor. For left ventricular ejection fraction, the optimal cutoff was EF = 51.7% (J = 0.775; Sensitivity = 0.900, Specificity = 0.875). For maximum heart rate during 24-hour Holter monitoring, the optimal cutoff was HRmax < 109 bpm (J = 0.600; Sensitivity = 0.800, Specificity = 0.800). The corrected cutoff was applied uniformly throughout all subsequent analyses (Firth logistic, Cox-Firth, and clinical score), with no re-optimization within the sensitivity framework, to preserve methodological uniformity across models.

#### 4.2 Two-Step Half-Up Rounding Rule for Score Weights

Integer weights of the clinical risk score were derived from the β coefficients of the retained multivariate model by a pre-specified two-step rounding rule applied uniformly to every predictor:

Step 1: Round each β coefficient to one decimal place using round-half-up (ties away from zero).

Step 2: Round the Step-1 result to the nearest integer using the same round-half-up rule.

This rule avoids asymmetric rounding between predictors and yields an integer β-ratio directly comparable across modeling frameworks. For the Firth logistic model: EF β = +4.481 → 4.5 → 5; HRmax β = +3.250 → 3.3 → 3; raw ratio 1.379 collapses to the irreducible integer ratio 5:3 → 2:1. For the Cox-Firth sensitivity model (Section 5): EF β = +2.317 → 2.3 → 2; HRmax β = +1.148 → 1.1 → 1; ratio 2.017 yields 2:1 directly. Both frameworks converge on the final weights {2, 1}, producing:

Score = 2 × [EF < 51.7%] + 1 × [HRmax < 109 bpm], range 0–3.

This convergence is the primary justification for adopting a single integer score applicable under either the logistic or the time-to-event interpretation of the data.

### 5. Cox-Firth Proportional Hazards Sensitivity Analysis

#### 5.1 Rationale

Follow-up duration differed systematically between survivors and nonsurvivors, and the Firth logistic model treats the 10-year mortality endpoint as a fixed binary outcome, ignoring time-to-event information. To address this asymmetry and to provide time-aware effect estimates, a Cox proportional hazards model with Firth-type penalization was fitted as a sensitivity analysis. The logistic regression remains the primary model, because the 10-year endpoint window and the echocardiographic assessment period used for Lyapunov exponent computation are temporally contemporaneous.

#### 5.2 Model Specification

The Cox-Firth model was fitted with the same two predictors retained by the primary logistic model — [EF < 51.7%] and [HRmax < 109 bpm] — without re-optimization of cutoffs. Implementation used the Python lifelines library (CoxPHFitter) with Firth-type penalization controlled by the penalizer parameter λ = 0.1. Time-to-event was defined as the interval in months from admission to either the endpoint event (nonsurvival) or administrative censoring at 10 years. Cutoffs were NOT re-optimized within the Cox framework, to preserve methodological uniformity with the primary analysis.

#### 5.3 Proportional Hazards Assumption

The proportional hazards assumption was assessed by Schoenfeld residuals with rank-transformed time, globally and per covariate. Both predictors satisfied the assumption: [EF < 51.7%] p = 0.696; [HRmax < 109 bpm] p = 0.334. No time-varying covariate or stratification correction was required.

#### 5.4 Discrimination and Optimism Correction

Model discrimination was quantified by Harrell’s concordance index (C-index). Optimism correction used bootstrap resampling (B = 1,000 resamples), following the same procedure adopted for the logistic AUC: for each resample, the model was refitted on the bootstrap sample, the optimism was computed as (C_boot_on_boot − C_boot_on_original), and the mean optimism was subtracted from the apparent C-index. Apparent C-index was 0.9344, and Cox optimism-corrected C-index was 0.9291 (95% CI 0.8856 – 0.9760), essentially identical to the logistic optimism-corrected AUC (0.959; 95% CI 0.923 – 1.007).

#### 5.5 Coefficient Stability

Bootstrap stability of the Cox-Firth β coefficients (B = 1,000) was as follows. EF < 51.7%: β = +2.376 ± 0.387, proportion β > 0 = 100.0%, proportion p < 0.05 = 99.5%. HRmax < 109 bpm: β = +1.152 ± 0.391, proportion β > 0 = 99.6%, proportion p < 0.05 = 51.8%. This hierarchy of robustness is concordant with the integer weights assigned to each predictor in the clinical score (2 for EF, 1 for HRmax).

#### 5.6 Convergence of Modeling Frameworks

The Firth logistic model and the Cox-Firth sensitivity model, despite differing in outcome formulation (binary vs time-to-event), converge on (i) the same two retained predictors, (ii) essentially identical bootstrap optimism-corrected discrimination (AUC 0.959 vs C-index 0.929), (iii) the same integer weight structure 2:1 under the two-step rounding rule (Section 4.2), and (iv) statistically significant independent effects of both predictors. This convergence supports the derivation of a single clinical risk score applicable under either modeling framework.

### 6. Logistic Firth Bootstrap Optimism Correction

The apparent AUC of the Firth logistic model was corrected for optimism using the Efron-Gong bootstrap procedure (B = 1,000 resamples). For each resample b: (i) the Firth model was refitted on the bootstrap sample; (ii) AUC_boot_on_boot was computed on the bootstrap sample; (iii) AUC_boot_on_original was computed on the full original sample using the bootstrap-fitted coefficients; (iv) optimism_b = AUC_boot_on_boot − AUC_boot_on_original. The mean optimism across 1,000 resamples was subtracted from the apparent AUC to yield the optimism-corrected estimate. Apparent AUC = 0.9650; mean optimism = 0.0064; optimism-corrected AUC = 0.9586, 95% CI [0.9228, 1.0072] (percentile method). Bootstrap stability of the Firth coefficients: EF < 51.7% β = +4.539 ± 0.712, proportion β > 0 = 100.0%, proportion p < 0.05 = 98.8%; HRmax < 109 bpm β = +3.308 ± 0.958, proportion β > 0 = 99.8%, proportion p < 0.05 = 92.4%.

### 7. Post-hoc Power and Sensitivity Analyses

**7.1 Background and rationale**

The present Supplementary Note reports post-hoc statistical power and sensitivity analyses for the two primary endpoints of the parent manuscript, aligned with the denominators used to derive the main-text statistical conclusions: (i) the Lyapunov exponent (LE) comparison between survivors and nonsurvivors in the cohort N = 50 (40 survivors / 10 nonsurvivors); and (ii) the Spearman rank correlation between the Study-Derived Score and the LE, also computed on the same N = 50 cohort, with a one-sided alternative hypothesis pre-specified in Methods (Statistical Analysis) (H₁: ρ < 0).

Two conceptually distinct post-hoc formulations are reported, with the recommendation of Greenland et al. and Lakens to prioritise the sensitivity-based interpretation (Approach 2). Observed power (Approach 1) in the same test from which a p-value was obtained is a monotone transformation of the p-value and conveys no incremental information beyond the p-value itself (Hoenig & Heisey, 2001); it is reported for methodological transparency only. The sensitivity-based minimum detectable effect size at α = 0.05 and power = 0.80 (Approach 2) characterises the design's detection floor independently of the observed statistic and provides the substantive power assessment.

**7.2 Methods**

**Primary analysis — LE survivors vs. nonsurvivors (cohort N = 50).** Source: Supplementary Table S1, row "Primary (N = 50)". Summary statistics: n1 = 40, mean LE = +0.349, SD = 0.342 (survivors); n2 = 10, mean LE = −0.459, SD = 0.377 (nonsurvivors). Pooled SD = 0.349. Observed Cohen's d = 2.32. For the two-sample t-test (equal variance, two-sided, α = 0.05), observed power was computed analytically using the noncentral t distribution with λ = d·√(n1·n2 / (n1 + n2)) and df = 48. For the Mann-Whitney U test, observed power was obtained by Monte-Carlo simulation (Bsim = 5 000) under a Gaussian shift model with the pooled SD as scale.

**Secondary analysis — Spearman Score–LE correlation (N = 50, one-sided).** Observed ρ = −0.369, p = 0.004 (one-sided), derived in the parent manuscript Results (Clinical Risk Score and Lyapunov Exponent). Observed power was computed analytically using the Fisher z-transformation: z(ρ) = arctanh(ρ), SE[z] = 1 / √(n − 3); one-sided power = Φ(|z(ρ)|·√(n − 3) − zα). Sensitivity: minimum detectable |ρ| was obtained by inverting the same relation for target power = 0.80.

**Bootstrap.** Parametric bootstrap with B = 1 000 resamples. For the LE analysis, each resample drew n1 and n2 Gaussian values from N(mi, SDi²) for each outcome group, recomputed d, and evaluated power at the resampled d. For the Spearman analysis, each resample drew n = 50 pairs from a bivariate Gaussian with the observed correlation structure, recomputed ρ and Fisher z, and evaluated one-sided power at the resampled |ρ|. Parametric bootstrap was chosen over nonparametric resampling for consistency with the low-n2 panels of Supplementary Figure S1 (panels E, L, M, P).

**Software.** Python 3 (scipy.stats, numpy). Noncentral-t via *scipy.stats.nct* with a numerical guard at |λ| > 7 (power saturates to 1.0). Mann-Whitney via *scipy.stats.mannwhitneyu* (two-sided, asymptotic with continuity correction). Full reproducible code included in the Supplementary Code archive.

**7.3 Results**

The power analysis for the primary LE comparison is summarised in Table S2 and for the Spearman correlation in Table S3.

**Table S2.** Post-hoc observed power (Approach 1) and sensitivity-based minimum detectable effect (Approach 2) for the primary LE comparison between survivors (n1 = 40) and nonsurvivors (n2 = 10) in the cohort N = 50. Observed Cohen’s d = 2.32.

| **Metric** | **t-test (primary)** | **Mann-Whitney U (secondary)** |
| --- | --- | --- |
| Observed effect size (Cohen's d) | 2.32 | 2.32 |
| Observed Δmean (LE units) | +0.809 | +0.809 |
| p-value (two-sided, as reported) | **< 0.001** | **< 0.001** |
| ***Approach 1 — Observed power (for completeness; Hoenig-Heisey caveat)*** | | |
| Analytical (noncentral t, guarded) | 1.000 | — |
| Simulation (B_sim = 5 000) | — | 1.000 |
| Parametric bootstrap mean ± SD (B = 1 000) | 0.998 ± 0.013 | — |
| Bootstrap 95 % percentile CI | [0.975, 1.000] | — |
| ***Approach 2 — Sensitivity: minimum detectable effect at α = 0.05, power = 0.80 (primary interpretation)*** | | |
| Minimum detectable Cohen's d | **1.01** | **1.05** |
| Equivalent minimum detectable Δmean (LE units) | +0.426 | +0.443 |

**Table S3.** Post-hoc observed power (Approach 1) and sensitivity-based minimum detectable effect (Approach 2) for the Spearman rank correlation between the Study-Derived Score and the Lyapunov exponent (N = 50, one-sided H₁: ρ < 0). Observed |ρ| = 0.381.

| **Metric** | **Spearman Score-LE (one-sided, H₁: ρ < 0)** |
| --- | --- |
| n | 50 |
| Observed ρ | −0.369 |
| p-value (one-sided, as reported) | **0.004** |
| ***Approach 1 — Observed power*** | |
| Analytical (Fisher z-transform, one-sided) | 0.844 |
| Parametric bootstrap mean ± SD (B = 1 000) | 0.722 ± 0.254 |
| Bootstrap 95 % percentile CI | [0.128, 0.998] |
| ***Approach 2 — Sensitivity (primary interpretation)*** | |
| Minimum detectable \|ρ\| at α = 0.05, power = 0.80 (one-sided) | **0.348** |

**7.4 Interpretation**

The primary LE comparison between survivors and nonsurvivors in the cohort N = 50 was robustly powered. The observed effect size (Cohen’s d = 2.32) was well above the sensitivity-based minimum detectable effect at conventional 80 % power for both tests (t-test d = 1.01; Mann-Whitney U d = 1.05). Observed power was effectively at saturation for the t-test and for the Mann-Whitney U. These values are consistent with the p-values reported in the parent manuscript (t-test p < 0.001; MW p < 0.001) and support the directional conclusion that survivors exhibit positive LE and nonsurvivors exhibit negative LE.

The secondary Spearman Score–LE correlation (|ρ| = 0.381, one-sided) was adequately powered at the design specification: the sensitivity-based minimum detectable |ρ| was 0.348, and the observed value exceeded this floor by a narrow but positive margin (Δ = 0.021). The analytical observed power of 0.844 is consistent with the reported one-sided p = 0.004. The bootstrap 95 % CI for |ρ| spanned [−0.58, −0.07], excluding zero in most resamples (observed power 95 % CI [0.128, 0.998]); the substantial lower boundary of the power CI reflects the fact that, at this sample size, a true correlation only modestly above the minimum detectable threshold admits a non-negligible tail of bootstrap resamples in which the effect is too small to reach significance.

***Caveat on imputation-induced variance compression.*** *The primary LE power analysis is performed on the cohort N = 50 because it matches the denominators of the main-text Spearman correlations and the LE comparison reported in Table 2 Panel A and the Results (Lyapunov Exponent and Mortality) of the parent manuscript. Because 32 of the 50 LE values are derived through outcome-stratified group-mean estimation of the missing derivatives, by construction the within-group variance is reduced relative to a hypothetical fully-measured design, and the corresponding Cohen’s d is modestly inflated. The sensitivity-based minimum detectable effect (Approach 2) is unaffected by this feature, as it depends only on n1, n2, α, and target power. Observed power (Approach 1) is therefore reported with a known optimistic bias, which reinforces the methodological preference for the sensitivity-based interpretation. Independent confirmation is provided by the restricted subset (n = 18; 16 survivors, 2 nonsurvivors): observed Cohen’s d = 2.02, sensitivity-based minimum detectable d = 2.22 (t-test) and 2.40 (MW), which, despite the adverse n2 = 2 configuration, places the observed d in the restricted subset marginally below the 80 %-power floor (a 9-16 % gap), consistent with the divergence between t-test p = 0.016 and MW p = 0.078 reported in that subset.*

***Hoenig-Heisey caveat.*** *Observed power (Approach 1) in the same test from which a p-value was obtained is a deterministic monotone transformation of the p-value and therefore does not constitute independent evidence. The Approach 1 values reported in Tables S1 and S2 are included for methodological transparency. The substantive conclusion of the present analysis relies on the sensitivity-based minimum detectable effect (Approach 2), which characterises the design independently of the observed statistic: for the primary endpoint (LE comparison, cohort N = 50), the design was adequately powered to detect effects of d ≥ 1.0, and the observed d = 2.32 comfortably exceeds this floor; for the secondary endpoint (Spearman Score–LE, N = 50), the design was adequately powered to detect correlations of |ρ| ≥ 0.35, and the observed |ρ| = 0.381 is just above this floor.*

### 8. Independent Validation in an Extended Follow-Up Sub-Cohort

**8.1 Rationale**

The primary analysis used the inter-patient finite-difference Lyapunov exponent (LE), motivated by the ergodic equivalence between inter-patient and intra-patient divergence under stationarity of the cohort dynamics (Methods, Section 5). Twelve patients of the same cohort had additional echocardiograms available after the initial follow-up, enabling independent estimation of the intra-patient LE.

**8.2 Methods**

LV mass at each time point was computed by the Devereux 1986 formula: LVM(g) = 1.04 × [(IVS + LVIDd + PWd)³ − LVIDd³] − 13.6, with measurements in cm. The intra-patient LE was estimated as the mean of log₁₀(|Δᵢ₊₁|/|Δᵢ|), where Δᵢ is the increment in LV mass between consecutive time points (3–5 measurements per patient; median extended observation 17.6 years beyond the original 10-year endpoint, range 10.4–19.8). Two versions were computed: (i) without temporal normalization; (ii) with normalization by the inter-examination interval, expressing the increments as rates (g/year). The Spearman correlation between intra-patient LE and each of the three clinical scores (Study Score, Rassi, Los Andes) was computed and validated by nonparametric bootstrap resampling (B = 1,000).

**8.3 Results**

Eleven of the 12 patients were 10-year survivors at the original endpoint; one died in 2020, beyond the original endpoint. The intra-patient LE (g/year) showed a robust negative correlation with the Rassi score: Spearman ρ = −0.662, one-sided p = 0.010; nonparametric bootstrap resampling (B = 1,000) confirmed the directional robustness, with ρ < 0 in 98.9 % of resamples (95 % CI [−0.925, −0.177]). The non-normalized version preserved the direction with marginal significance (ρ = −0.473, p = 0.060). Correlations with the Study Score and Los Andes did not reach significance, reflecting the low variance of these scores within the sub-cohort (Study Score: 10 of 12 at Score 0; Los Andes: 9 of 12 at stage II). Per-patient values are reported in Supplementary Table S4.

**8.4 Significance**

The intra-patient LE recovered the same direction and similar magnitude as the inter-patient LE Rassi gradient (inter-patient ρ = −0.243, N = 50; intra-patient ρ = −0.662, n = 12), despite operating at a different temporal scale and on an independent set of derivatives. The convergence supports the ergodic equivalence assumed in the primary methodology. Limitations include the small sub-cohort, its predominance of survivors, the absence of variation in the Study Score, and the heterogeneous inter-examination intervals, which the rate-normalized LE was designed to address.

**Supplementary Table S4.** Per-patient intra-patient Lyapunov exponent (LE) in the extended follow-up sub-cohort (n = 12). LE was computed as the mean of log₁₀(|Δᵢ₊₁|/|Δᵢ|) across 3–5 longitudinal LV mass measurements per patient. LE (no Δt) uses unnormalized increments; LE (g/year) uses time-normalized increments. Bold row identifies the patient with documented death after the original 10-year endpoint.

| **PID** | **n pts** | **LE (no Δt)** | **LE (g/year)** | **Study Score** | **Rassi** | **Los Andes** | **Outcome** |
| --- | --- | --- | --- | --- | --- | --- | --- |
| P001 | 5 | +0.079 | +0.724 | 0 | 2 | 2 | alive |
| P003 | 4 | +0.108 | +0.063 | 0 | 0 | 2 | alive |
| P016 | 4 | -0.348 | +0.513 | 0 | 0 | 2 | alive |
| P018 | 4 | +0.140 | +0.073 | 0 | 3 | 2 | alive |
| P021 | 3 | -0.003 | -0.485 | 0 | 5 | 2 | alive |
| P025 | 4 | +0.622 | +0.739 | 0 | 0 | 2 | alive |
| P027 | 3 | +0.321 | +1.057 | 0 | 0 | 1 | alive |
| P031 | 3 | -0.551 | -0.790 | 0 | 2 | 1 | alive |
| **P032** | **3** | **-0.009** | **-0.349** | **0** | **2** | **2** | **died 2020-10** |
| P033 | 3 | -0.458 | -0.484 | 0 | 3 | 2 | alive |
| P036 | 3 | -0.607 | -1.085 | 1 | 5 | 3 | alive |
| P040 | 5 | +0.462 | +0.853 | 1 | 2 | 2 | alive |
